# Clonal Hematopoiesis Instructs Maladaptive Tissue Repair to Promote Fibrosis

**DOI:** 10.64898/2026.03.24.710438

**Authors:** Dongzhu Li, Ana C. Viñado, Paula Garcia-Olloqui, Sydney B. Montesi, Barry S. Shea, Leonard Christian, Iria Vazquez-Urio, Paula Aguirre-Ruiz, Beñat Ariceta, Manuela Neubert, Ashutosh Tripathy, Niloy Barua, Pablo S. Valera, Laura Vera, Eva Petri, Antje Prasse, Benjamin Seeliger, Amaia Vilas, Patxi San Martin, Maria José Calasanz, Felipe Prosper, Peter G. Miller, Werner Seeger, Florian H. Heidel, Jonas C. Schupp, Borja Saez, Ana Pardo-Saganta

**Affiliations:** Institute for Lung Health (ILH), Justus-Liebig University, Universities of Giessen and Marburg Lung Center (UGMLC), German Center for Lung Research (DZL), Giessen, Germany; Cardio-Pulmonary Institute (CPI), Department of Internal Medicine, Justus Liebig University; Hematology and Oncology Program, Centre for Applied Medical Research (CIMA), Instituto de Investigaciones Sanitarias de Navarra (IdiSNA), Cancer Center Clinica Universidad de Navarra (CCUN), Pamplona, Navarra, Spain; Division of Pulmonary and Critical Care Medicine, Massachusetts General Hospital, Boston, Massachusetts; Department of Respiratory Medicine and Infectious Diseases, Hannover Medical School, Hannover, Germany; Biomedical Research in End-Stage and Obstructive Lung Disease (BREATH), Hannover Medical School (MHH), German Center for Lung Research (DZL), Hannover, Germany; CIMA LAB Diagnostics-Clínica Universidad de Navarra, Pamplona, Navarra Institute for Health Research (IdiSNA), Spain; Innere Medizin C, Universitätsmedizin Greifswald, Greifswald, Germany; Fraunhofer Institute for Toxicology and Experimental Medicine, Hannover, Germany; Centro de Investigacion Biomedica en Red de Cancer (CIBERONC), Madrid, Spain; Hematology and Cell Therapy Department. Cancer Center Clinica Universidad de Navarra (CCUN), IdiSNA, Pamplona, Spain; Center for Regenerative Medicine and Krantz Family Center for Cancer Research, Massachusetts General Hospital, Boston, MA, USA; Broad Institute of the Massachusetts Institute of Technology and Harvard University, Cambridge, MA, USA; Harvard Medical School, Boston, MA, USA; Department of Internal Medicine, Universities of Giessen and Marburg Lung Center (UGMLC), Member of the German Center for Lung Research (DZL), Giessen, Germany; Institute for Lung Health (ILH), Cardio-Pulmonary Institute (CPI), Giessen, Germany; Department of Hematology, Hemostasis, Oncology and Stem Cell Transplantation, Hannover Medical School (MHH), Hannover, Germany; Leibniz Institute on Aging, Fritz-Lipmann-Institute, Jena, Germany; Section of Pulmonary, Critical Care, and Sleep Medicine, Department of Internal Medicine, Yale School of Medicine, New Haven, CT, USA

## Abstract

Tissue repair is increasingly recognized as a systemic process influenced by age-associated changes beyond the injured organ itself. Clonal hematopoiesis of indeterminate potential (CHIP), a common consequence of somatic evolution in hematopoietic stem cells, has been linked to inflammatory disorders, yet whether it directly regulates tissue remodeling remains unclear.

Here, we integrate population genomics, preclinical models, and human lung analyses to examine the role of CHIP in fibrotic lung disease. In large cohorts, idiopathic pulmonary fibrosis (IPF) was associated with a distinct CHIP mutational spectrum enriched for non-*DNMT3A* variants and for larger mutant clones. In mouse models, hematopoietic mutations exacerbated bleomycin-induced fibrosis and reprogrammed macrophages toward inflammatory, profibrotic states, including expansion of a distinct, injury-responsive SPP1^+^ population conserved in human disease. CHIP-associated macrophages were sufficient to directly promote fibroblast activation and alter epithelial differentiation, linking hematopoietic genotype to parenchymal remodeling. Consistently, a CHIP-derived macrophage transcriptional signature predicted adverse outcomes in independent IPF cohorts. Notably, immune and epithelial alterations were detectable even in the absence of overt injury, indicating that CHIP establishes a primed tissue environment permissive for maladaptive repair.

Together, these findings identify clonal hematopoiesis as a systemic regulator of tissue repair and demonstrate that somatic evolution in blood can actively instruct organ remodeling through immune-parenchymal interactions. This framework supports the possibility that disease-associated selective pressures may shape clonal architecture with functional consequences for organ health.

## Introduction

Idiopathic pulmonary fibrosis (IPF) is a progressive, fatal interstitial lung disease that predominantly affects older adults, characterized by irreversible scarring of the lung parenchyma, impaired gas exchange, and median survival of fewer than five years after diagnosis^1–3^. Although antifibrotic therapies modestly slow disease progression^4–6^, IPF remains incurable, and substantial heterogeneity in disease course and therapeutic response persists^1–3^. Moreover, the absence of clinically actionable biomarkers that capture systemic drivers of disease, contributes to the inability to predict progression, stratify patients, or tailor therapeutic strategies in IPF. While lung-specific processes such as epithelial injury and aberrant repair are central to disease pathogenesis^7^, the consistent association with advanced age suggests that systemic factors also shape susceptibility to lung fibrosis. This perspective reframes IPF as a manifestation of organismal aging, in which age-dependent changes in distant organs or tissues may prime the lung for maladaptive remodeling.

Emerging evidence suggests that aging-related changes in the hematopoietic system act as a hidden driver of systemic vulnerability^8,9^. Hematopoietic stem cells accumulate somatic mutations and undergo clonal expansion over time, generating persistent low-grade local and systemic inflammation ^10,11^. Clonal hematopoiesis of indeterminate potential (CHIP), the age-associated presence of mutated hematopoietic clones, exemplifies this process^12^. CHIP clones act as enduring sources of proinflammatory and profibrotic signals, amplifying systemic stress without direct evidence of them causing organ disease^13^. While initially considered a benign marker of aging, evidence from multiple age-related pathologies demonstrates how CHIP clones amplify age-dependent susceptibility, largely through mutation-driven reprogramming of innate immune cells toward heightened inflammatory states^14–17^. These findings have reframed CHIP as an active modifier of chronic disease biology rather than a passive byproduct of hematopoietic aging^18^, raising the possibility that similar systemic mechanisms contribute to pulmonary fibrosis.

Despite the conceptual alignment between CHIP biology and IPF pathogenesis, the role of clonal hematopoiesis in pulmonary fibrosis has received limited attention. Prior studies have linked CHIP to chronic obstructive pulmonary disease (COPD) and increased susceptibility to pulmonary infections^19–22^, yet whether similar systemic influences shape susceptibility to fibrotic lung disease has not been systematically explored.

This gap is particularly notable given the central role of monocyte-macrophage lineages in fibrogenesis^23–25^. Macrophages are increasingly recognized as critical regulators of fibroblast activation, extracellular matrix deposition, and epithelial fate decisions in pulmonary fibrosis^26–28^, and represent a major cellular compartment shaped by CHIP-associated mutations^14,15,29^. Moreover, IPF presents a biological context in which systemic and local processes are likely to intersect. The lung is continuously exposed to environmental injury and oxidative stress, particularly in older individuals with smoking histories, raising the possibility that chronic lung disease itself may influence hematopoietic clonal dynamics.

Here, we investigated whether clonal hematopoiesis contributes to susceptibility, progression, and tissue remodeling in idiopathic pulmonary fibrosis. Leveraging whole-genome sequencing from large population-based cohorts, we examined the prevalence, mutational composition, clonal features and association of CHIP and IPF. Preclinical models of clonal hematopoiesis were combined with experimental analyses of murine and human lung tissues to define how CHIP-associated immune cells shape inflammatory responses, parenchymal remodeling, and fibrotic outcomes. Together, these approaches connect clonal architecture with immune reprogramming, tissue-level responses, and adverse clinical outcomes in fibrotic lung disease.

## Results

### CHIP is associated with IPF, with the association modulated by mutation identity and clone size

To determine the relationship between CHIP and IPF, we analyzed whole-genome sequencing data from 1211 individuals with IPF enrolled in the TOPMed Pulmonary Fibrosis Whole Genome Sequencing cohort (WGS-IPF). In addition, 2897 population-based participants from the TOPMed ARIC study (NHGRI CCDG: Atherosclerosis Risk in Communities) served^8,9,30^ as external controls due to the absence of controls within the IPF dataset. As expected, individuals with IPF were older than controls (mean age 65 ± 10 years vs. 58 ± 7.2 years; p<0.001), more often male (66% vs. 51%; p<0.001), and more frequently reported a history of smoking (69% vs. 51%; p<0.001) **(Extended Data Fig. 1A)**.

**Figure 1:**
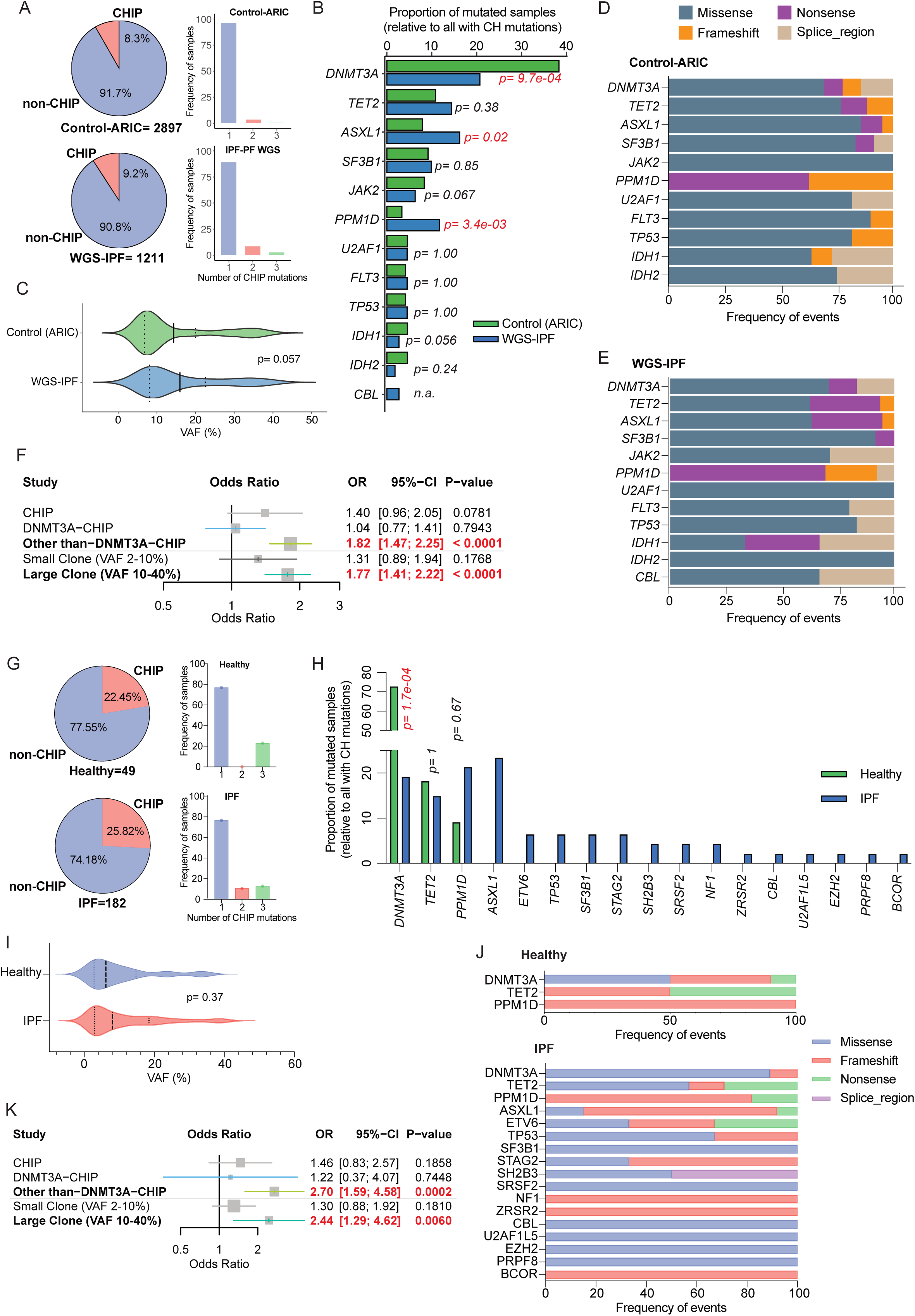
Mutation identity and clonal expansion define the association between CHIP and IPF. **a,** Prevalence of CHIP and number of variants per sample in two public TOPMed WGS cohorts: ARIC (control, n= 2897) and PFWGS (IPF, n= 1211). **b,** Relative abundance of CHIP-associated gene variants among all CHIP mutations identified in the WGS cohorts. **c,** Distribution of clone sizes, measured by variant allele fractions (VAFs), for CHIP mutations across WGS cohorts. **d,e,** Distribution of functional mutation types across CHIP variants in WGS cohorts. **f,** Logistic regression model, adjusted for sex, age, and smoking status, showing the association between CHIP (and CHIP-other-than-DNMT3A) and IPF, and the effect of clone size (large clone: VAF 10-40%; small clone: VAF 2-10%) in the WGS cohorts. **g,** Prevalence of CHIP and number of variants per sample in an independent validation cohort (n= 49 healthy controls; n= 182 IPF patients) assessed by targeted DNA sequencing. **h,** Relative abundance of CHIP-associated gene variants among all CHIP mutations in the validation cohort. **i,** Distribution of clone sizes (VAFs) for CHIP mutations in the validation cohort. **j,** Distribution of functional mutation types across CHIP variants in the validation cohort. **k,** Logistic regression model, adjusted for sex, age, and smoking status, showing the association between CHIP (and CHIP-other-than-DNMT3A) and IPF, and the effect of clone size (large clone: VAF 10–40%; small clone: VAF 2–10%) in the validation cohort. Results in **f** and **k** are presented as odds ratios (ORs) with 95% confidence intervals and p-values.

We identified CHIP mutations in 240 (8.3%) of the 2897 controls from the ARIC study and in 111 (9.2%) of the 1211 cases from the WGS-IPF study **(Fig. 1A)**. The number of mutations per individual was comparable across groups **(Fig. 1A)**. Among controls, the most frequently mutated genes were *DNMT3A*, *TET2*, and *ASXL1* **(Fig. 1B)**, consistent with established CHIP patterns^9,31^. In contrast, IPF cases demonstrated a higher incidence of *ASXL1* and *PPM1D* mutations and a lower frequency of *DNMT3A* mutations, while the prevalence of *TET2* mutations remained comparable between groups **(Fig. 1B)**. These differences in the prevalence of specific CHIP mutations between IPF cases and controls were consistent across age groups, sexes, and smoking-status categories **(Extended Data Fig. 1B-D)**. Among CHIP carriers, the median variant allele fraction (VAF) **(Fig.1C)** and the prevalence of large clones (VAF ≥10%) was comparable between IPF cases and controls (55% vs. 45%; p<0.11) **(Extended Data Fig. 1A)**. Clone size increased with age similarly in both IPF and control subjects **(Extended Data Fig. 1E)**. Functional mutation classes were comparable across cohorts and consistent with their previously described distribution for CHIP **(Fig 1D,E)**^8,9,30^.

In a multivariable logistic regression model adjusting for age, sex, and smoking status, overall CHIP was not significantly associated with IPF (adjusted odds ratio [OR] 1.40; 95% CI 0.96-2.05; p= 0.0781) **(Fig. 1F)**. Given the markedly lower prevalence of *DNMT3A* mutations in IPF cases, we next examined the association of CHIP excluding *DNMT3A* variants. Notably, CHIP driven by non-*DNMT3A* mutations was associated with increased odds of IPF (adjusted OR 1.82; 95% CI 1.47-2.25; p< 0.0001) **(Fig. 1F)**. Clone size further modified this relationship; individuals with large clones (VAF ≥10%), including *DNMT3A*, had higher odds of IPF (adjusted OR 1.77; 95% CI 1.41-2.22; p< 0.0001), whereas smaller clones showed no significant association (adjusted OR 1.31; 95% CI 0.89-1.94; p= 0.1768) **(Fig. 1F)**.

To validate these observations, we sequenced an independent cohort of IPF patients (n= 182) and healthy controls (n= 49) from two institutions using targeted DNA sequencing panels covering all CHIP-associated genes. Patients’ characteristics are summarized in (**Extended Data Fig. 1F)**. We identified CHIP mutations in 11 (22.45%) of the 49 controls and in 47 (25.82%) of the 182 IPF cases, with a similar number of mutations per individual in both groups **(Fig. 1G)**. Consistent with the observations in the larger WGS-IPF and ARIC datasets, IPF cases in the validation cohort displayed the same distinct gene-specific CHIP distribution in IPF compared to control individuals **(Fig. 1H)**. These differences were stable across age, sex, and smoking-status strata in our validation cohort **(Extended Data Fig. 1G-I).** This disease specific enrichment in CHIP variants was further corroborated in a third independent dataset (LTRC_IPF) **(Extended Data Fig. 1J)**. Further, clone sizes and their increase with age, prevalence of large clones and functional mutation distributions remained comparable **(Fig.1I,J and Extended Data Fig. 1K).**

Logistic regression in the validation cohort recapitulated the GWAS findings, showing that other-than-*DNMT3A* CHIP was associated with increased odds of IPF (adjusted OR 2.70; 95% CI 1.59-4.58; p= 0.0002) and that larger clones (including *DNMT3A*) were similarly linked to higher odds of disease (adjusted OR 2.44; 95% CI 1.29-4.62; p= 0.006) **(Fig.1K)**.

These findings suggest that both the identity of the mutated gene and the extent of clonal expansion influence the relationship between CHIP and IPF, consistent with a potential role of hematopoietic somatic mutations in modulating systemic inflammation or tissue remodeling relevant to pulmonary fibrosis.

### Mendelian randomization supports a gene- and clone-size–dependent causal contribution of CHIP to IPF

To assess potential causal contributions of CHIP to IPF, we performed two-sample Mendelian randomization (MR) using genetic instruments derived from two large population-based datasets^32^, followed by meta-analysis. Genetically predicted CHIP was positively associated with IPF risk, with the strongest causal estimates observed for *TET2* and *PPM1D* mutations, whereas *DNMT3A and ASXL1*-driven CHIP showed weaker effects **(Fig. 2A)**. Importantly, larger CHIP clones amplified the causal effect of CHIP to IPF risk **(Fig. 2A)**, highlighting clone size as a key modifier of CHIP’s impact.

**Figure 2:**
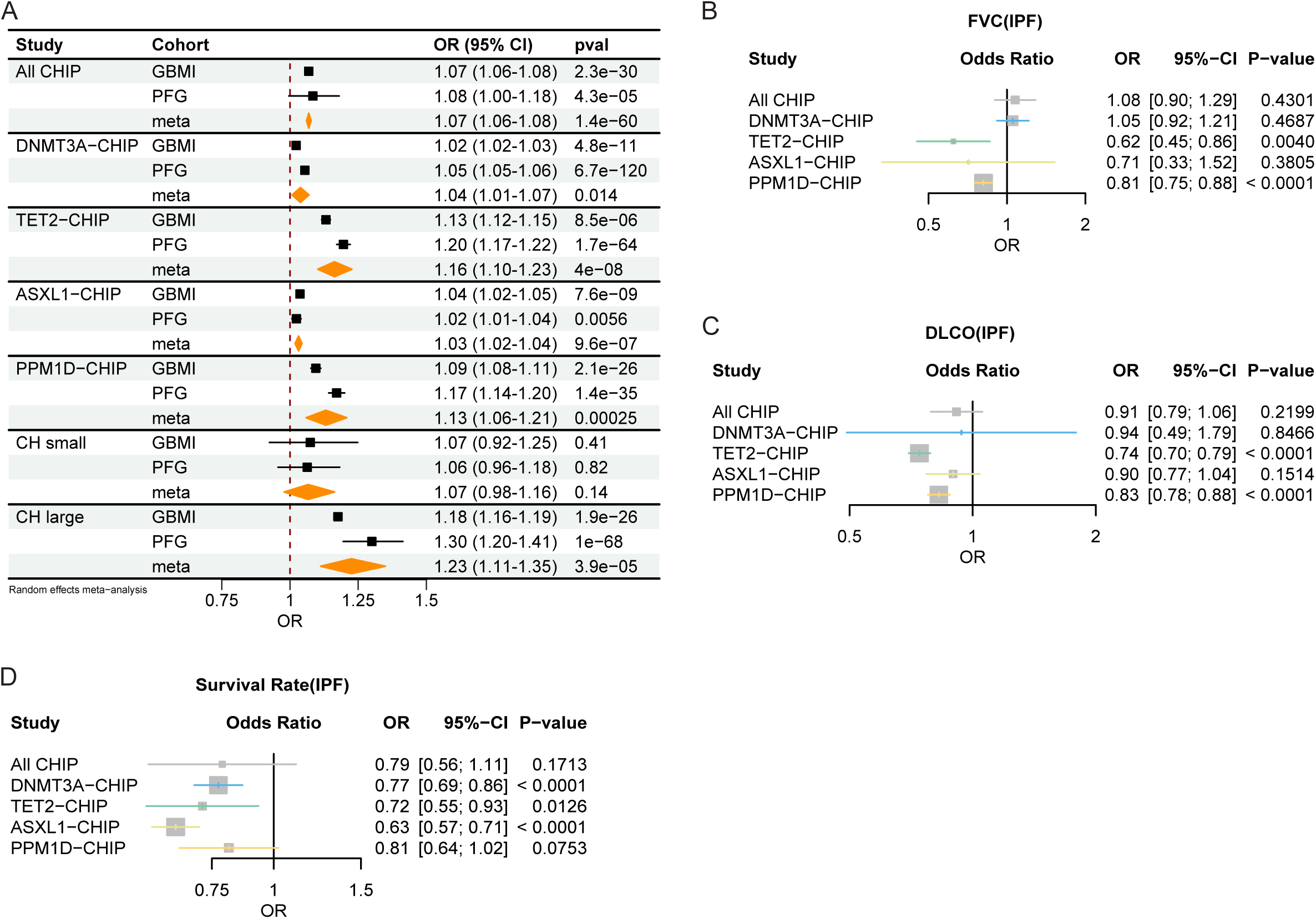
Mendelian randomization supports a gene- and clone-size–dependent causal contribution of CHIP to IPF. **a,** Mendelian randomization (MR) assessing the association between CHIP and IPF risk. Results represent a random-effects meta-analysis using conventional multiplicative inverse-variance weighted (IVW) estimators, combining data from the GBMI and PFG cohorts. **b-d,** MR exploring the association between CHIP and clinical outcomes within the PFG cohort: **b,** Forced Vital Capacity (FVC), **c,** Diffusing Capacity of the Lung for Carbon Monoxide (DLCO), and **d,** Transplant-free Survival Rate. Results are presented as odds ratios (ORs) with 95% CIs and p-values.

Importantly, MR analyses revealed associations between CHIP and adverse clinical outcomes in IPF. Specifically, CHIP was linked to reduced lung function, including lower FVC (Forced Vital Capacity) and DLCO (Diffusing Capacity for Carbon Monoxide), with the magnitude of these associations varying by the underlying mutation **(Fig. 2B-C)**. In addition, CHIP was also associated with decreased survival, again with different gene mutations contributing unequally to causal estimates **(Fig. 2D)**. Notably, larger clones, considered across all CHIP mutations collectively, strengthened these associations for all outcomes **(Extended Data Fig. 2A-C)**. Conversely, reverse MR analyses testing the effect of IPF on CHIP showed no significant association **(Extended Data Fig. 2D)**. Together, these results indicate that both mutation identity and clonal expansion modify the impact of CHIP on IPF risk and progression, supporting a potential causal role in disease severity and clinical outcomes.

### Clonal hematopoiesis confers enhanced vulnerability to fibrotic remodeling after bleomycin injury

Having established the association between CHIP and IPF, as well as its impact on clinical outcomes, we next investigated its causal contribution to pulmonary fibrosis using experimental mouse models of clonal hematopoiesis.

We prioritized *Tet2* and *Asxl1* models since *DNMT3A* mutations were markedly decreased in IPF while *ASXL1* variants were selectively enriched, and *TET2* mutation frequencies were preserved across cohorts. Moreover, Mendelian randomization analyses identified both *TET2* and *ASXL1* as key contributors to the CHIP-related causal effects on IPF.

To model clonal hematopoiesis in mice, we performed bone marrow (BM) transplantation using *Tet2* or *Asxl1* wild-type (WT) or deficient bone marrow (BM) cells (from Tet2^fl/fl^-Mx1cre^−^, Tet2^fl/fl^-Mx1-cre^+^ or Asxl1^fl/fl^-Mx1cre^−^, Asxl1^fl/fl^-Mx1-cre^+^ mice), all on a C57BL/6J CD45.2 background (all donor mice received pIpC 10-15 days before BM harvest). These cells were mixed at a 1:9 ratio with WT BM from syngeneic B6-SJL donors (CD45.1) and transplanted into lethally irradiated recipients **(Extended Data Fig. 3A)**^15^. For clarity, mice receiving WT *Tet2* or *Asxl1* BM together with B6-SJL BM are referred to as Control no-CHIP mice, whereas those receiving *Tet2*- or *Asxl1*-deficient BM are referred to as *Tet2-*CHIP or *Asxl1-*CHIP mice, respectively. Nine weeks after BM transplantation, Control no-CHIP, Tet2-CHIP, and Asxl1-CHIP mice received a single dose of bleomycin to induce pulmonary fibrosis. At this time point, all groups exhibited robust hematopoietic engraftment **(Extended Data Fig. 3B)** with no notable differences in peripheral blood WBC, RBC, or platelet counts, and with comparable myeloid and lymphoid lineage distributions **(Extended Data Fig. 3C-E)**. As anticipated, *Tet2*-CHIP mice demonstrated progressive clonal expansion, consistent with the known hyperproliferative behavior of *Tet2*-deficient hematopoietic cells^33^, whereas clone size remained stable in *Asxl1*-CHIP mice throughout the experiment **(Extended Data Fig. 3B)**^34^. Fourteen days after bleomycin administration, *Tet2*-CHIP mice exhibited markedly more severe fibrotic pathology than Control no-CHIP mice, as evidenced by histological assessment **(Fig. 3A,B)**. *Tet2*-CHIP lungs showed a significant expansion of areas with moderate to severe tissue damage **(Fig. 3C)**, accompanied by increased collagen deposition reflected both by a larger collagen-positive area on Masson’s trichrome staining **(Fig 3. B,D)** and by enhanced Col1a1 immunofluorescence coverage **(Extended Data Fig. 3F,G)**. Consistent with these findings, hydroxyproline quantification, a biochemical surrogate of collagen accumulation, confirmed the heightened fibrotic burden in *Tet2*-CHIP mice **(Fig. 3E)**. *Asxl1*-CHIP mice showed a similar pattern of enhanced fibrosis, including greater tissue damage (Fig 3. F-H) and collagen deposition **(Fig. 3 I,J and Extended Data Fig. 3 H,I)**, relative to Control no-CHIP mice. Altogether, these results demonstrate that clonal hematopoiesis driven by *Tet2* or *Asxl1* amplifies the fibrotic cascade that follows lung damage.

**Figure 3:**
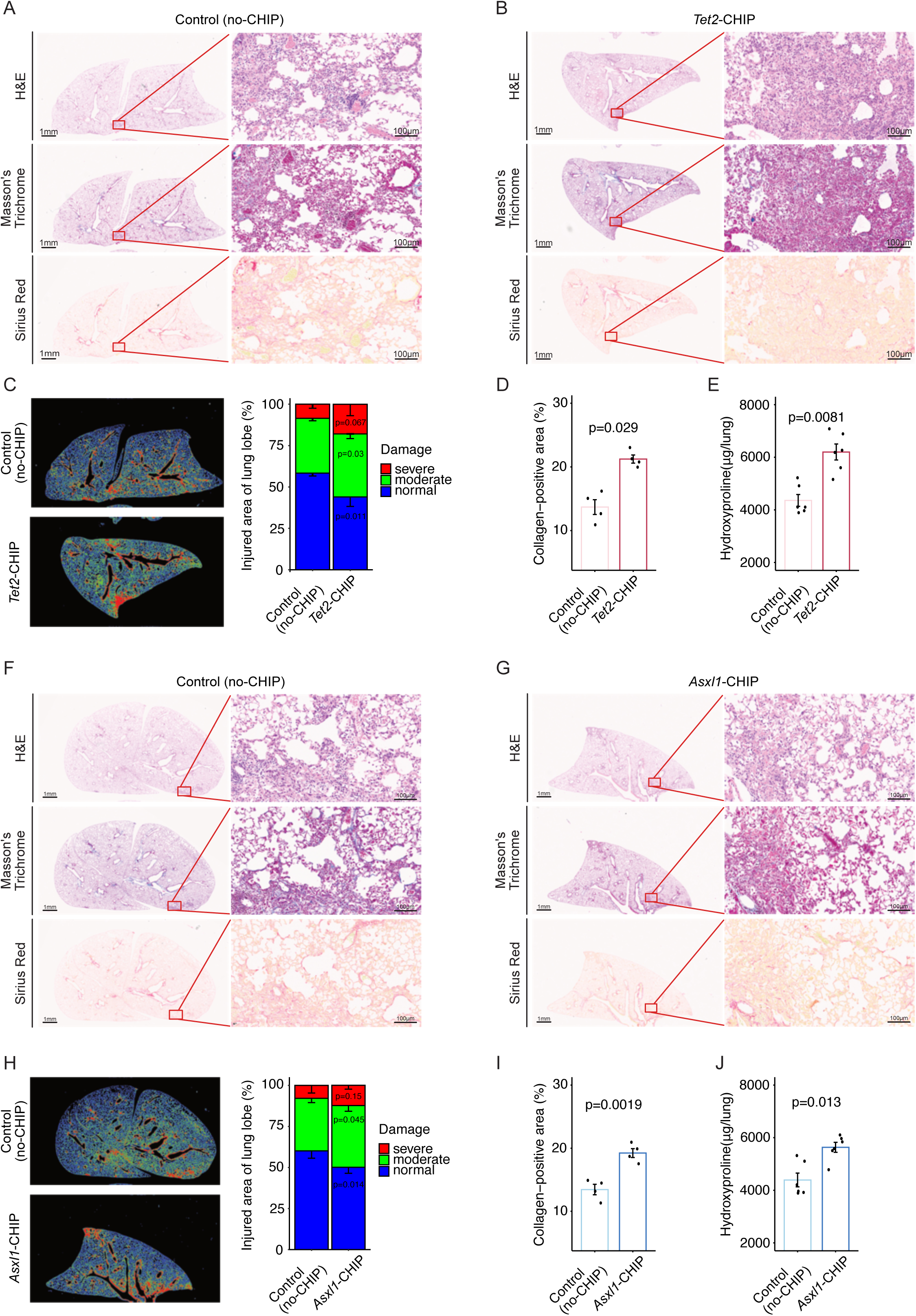
Clonal hematopoiesis confers enhanced vulnerability to fibrotic remodeling after bleomycin injury. **a,b,** Representative lung histology images from control mice (a) and *Tet2*-CHIP mice (b) 14 days post–bleomycin injury (14 d.p.i.), stained with Hematoxylin and Eosin (H&E), Masson’s Trichrome (collagen in blue), and Sirius Red. **c,** Lung injury severity in *Tet2*-CHIP mice assessed by automated image analysis. Left: representative segmentation showing normal (blue), moderately damaged (green), and severely damaged (red) regions. Right: quantification of injury severity expressed as the percentage of total lung area. **d,** Collagen area quantified from Masson’s Trichrome staining in *Tet2*-CHIP mice and matched controls. **e,** Hydroxyproline content in total lung homogenates at 14 d.p.i., providing a biochemical measure of total collagen deposition in *Tet2*-CHIP mice and matched controls. **f,g,** Representative lung histology images from control mice (left) and *Asxl1*-CHIP mice (right) 14 d.p.i., stained with H&E, Masson’s Trichrome, and Sirius Red. **h,** Lung injury severity in *Asxl1*-CHIP mice assessed by automated image analysis. Left: segmentation showing normal (blue), moderately damaged (green), and severely damaged (red) regions. Right: quantification of injury severity as the percentage of total lung area. **i,** Collagen area quantified from Masson’s Trichrome staining in *Asxl1*-CHIP mice and matched controls. **j,** Hydroxyproline content in total lung homogenates at 14 d.p.i., measuring total collagen deposition in *Asxl1*-CHIP mice and matched controls. Results shown represent independent experiments. n= 4-6 mice per genotype per experiment unless otherwise indicated. Data are presented as mean and standard error of the mean (SEM). Scale bars, 1 mm (low-magnification images) and 100 μm (high-magnification images).

### CHIP amplifies inflammatory macrophage programs and expands an injury-responsive SPP1^+^ macrophage population in fibrotic lung injury

Consistent with the amplified tissue injury and fibrosis observed in both CHIP models, cytokine profiling of lung lysates revealed that clonal hematopoiesis markedly intensifies the inflammatory response to bleomycin injury. Despite mutation-specific differences in *Tet2*- and *Asxl1*-CHIP mice, both models converged on a shared inflammatory program characterized by increased IL-1 family cytokines, TNF/TNFR-superfamily signaling, CXCL1-mediated neutrophil recruitment, CCL2/CCL5 chemotaxis pathways, and elevated TIMP-1 **(Fig. 4A)**.

**Figure 4:**
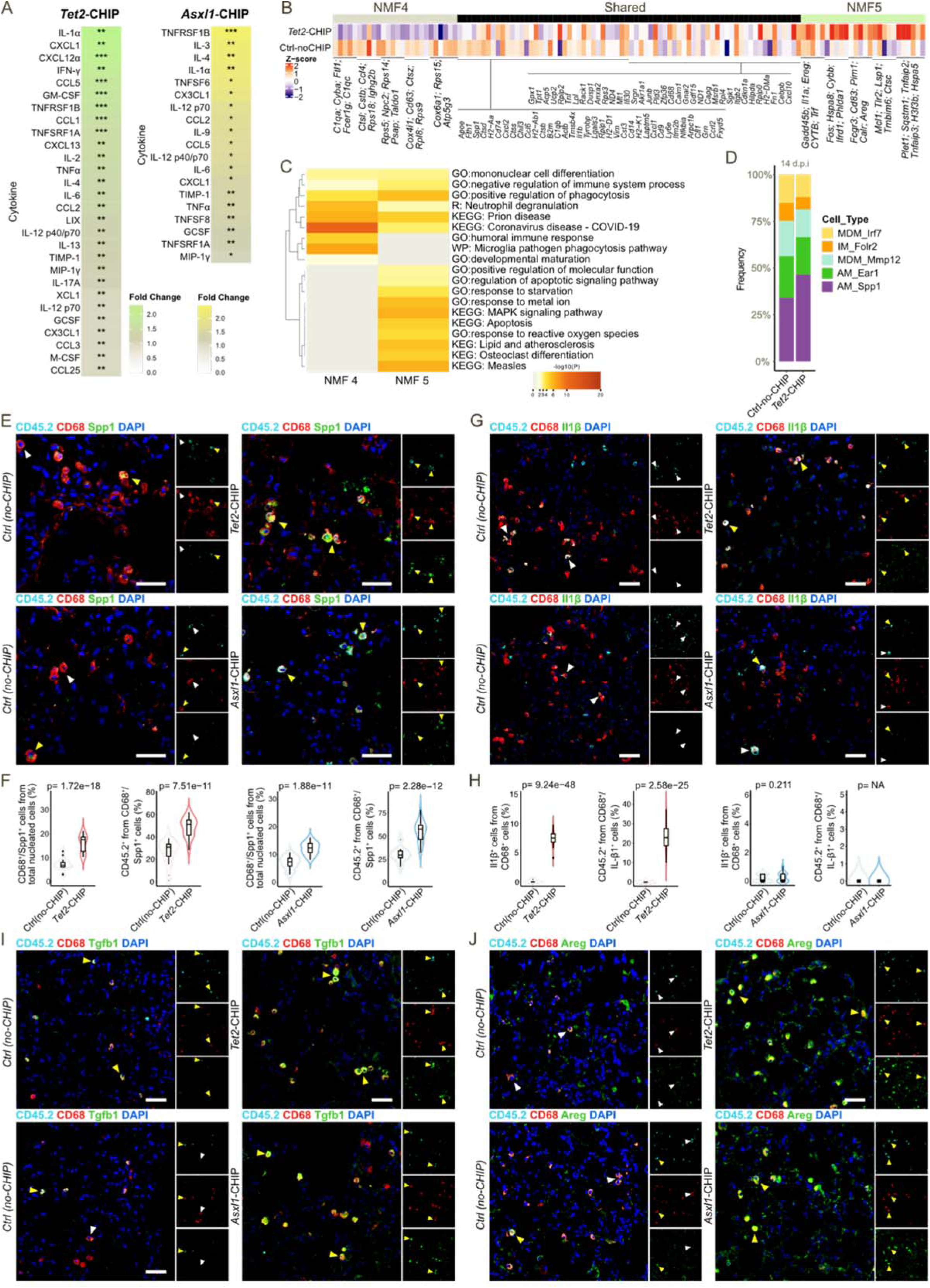
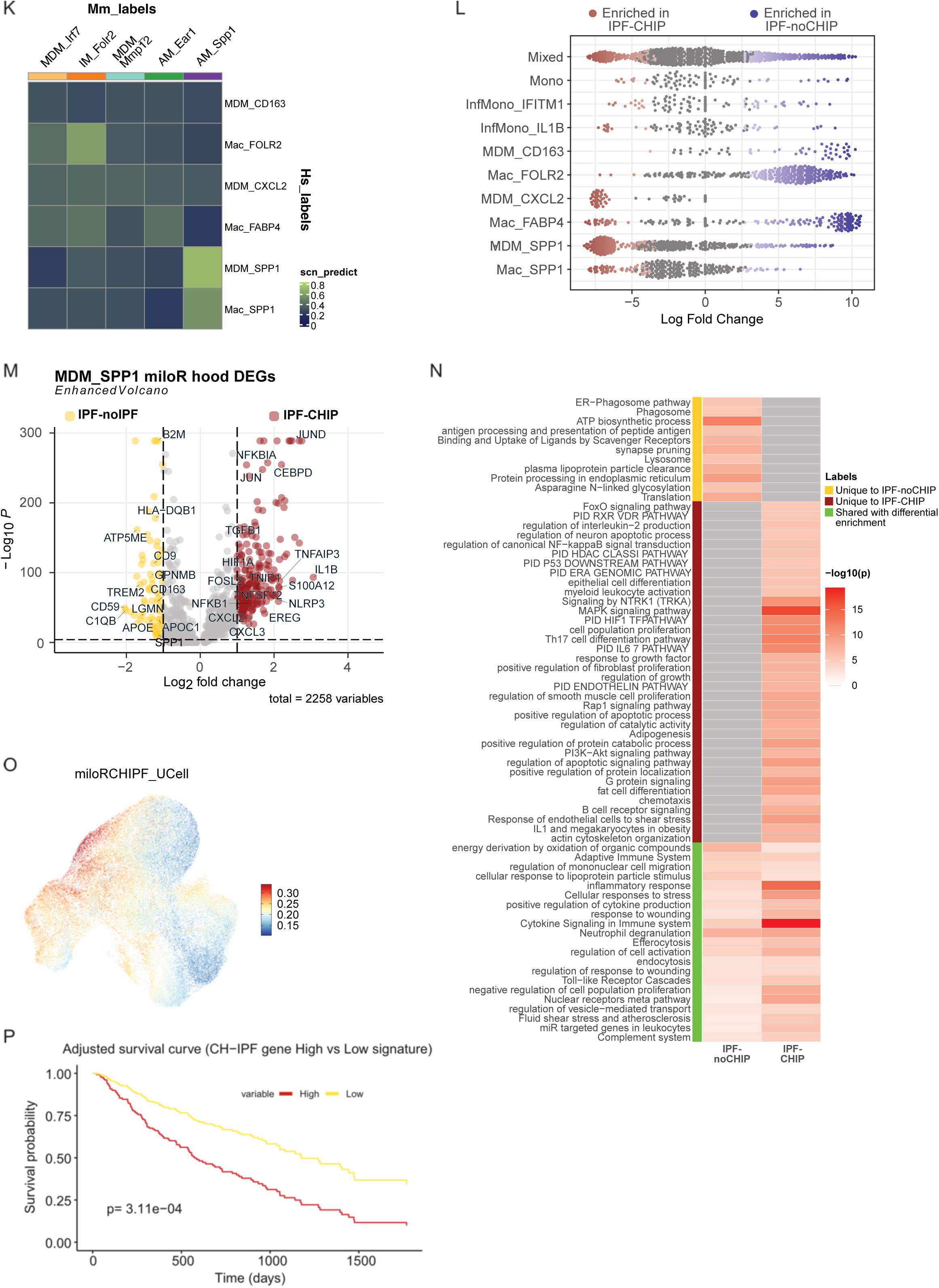
CHIP amplifies inflammatory macrophage programs and expands an injury-responsive Spp1^+^ population in fibrotic lung injury. **a,** Inflammatory cytokine profile in total lung homogenates from the *Tet2-* and *Asxl1-*CHIP mice compared to control-noCHIP mice at 14 d.p.i. showing fold change and p-val. **b,** Gene expression Z-scores of unique and shared genes associated with Non-negative Matrix Factorization (NMF) clusters 4 and 5 (NMF4, NMF5) derived from bulk RNA-seq data comparing control versus *Tet2*-CHIP mouse lung macrophages at 14 d.p.i. **c,** Comparative functional enrichment analysis of gene signatures enriched in NMF4 versus NMF5 in Control-noCHIP and *Tet2-*CHIP mice. **d,** Deconvolution analysis of the bulk RNA-seq data, estimating the percentage of different macrophage sub-clusters based on mouse macrophage scRNA-seq reference dataset (GSE280003) in Control-noCHIP and *Tet2-*CHIP mice 14 d.p.i. **e-j,** Representative immunofluorescence images and their quantification (when specified) of the co-localization of CD68^+^ and CD45.2^+^ macrophages with: e,f, Secreted Phosphoprotein 1 (SPP1); g,h, Interleukin-1 beta (IL-1beta); i, Transforming Growth Factor beta 1 (TGF beta 1) and j, Amphiregulin (AREG) in the lungs of *Tet2-* and *Asxl1-*CHIP mice at 14 d.p.i. Yellow arrowheads point to triple positive cells. White arrowheads point to CD68^+^ CD45.2^+^ cells that are negative for the third marker. **k,** Classification of similarity of single-cell RNA-Seq data across human (GSE227136) and mouse (GSE280003) macrophage datasets. **l,** Beeswarm plot of the log2(fold change) in the Milo neighborhoods between human IPF patients with and without predicted CHIP mutations, grouped by cell type. Neighborhoods showing a significant change in cellular abundance are colored. **m**, Volcano plot showing differentially expressed genes (DEGs) between IPF with and without predicted CHIP mutation within the MDM_SPP1 cluster, restricted to cells from Milo neighborhoods identified as significant in panel **l**. Genes with P-value <0.01 and log2(fold change) > 1 are color-coded. **n,** Comparative functional enrichment analysis of the gene signatures identified in panel **m** between IPF with and without predicted CHIP mutations. **o,** Module score of significant DEGs identified in panel **m**, representing the IPF with CHIP gene signature, visualized on the UMAP. **p.** Cox model-based adjusted survival curves for IPF patients stratified by expression of the CHIP-associated macrophage transcriptional signature, with adjustment for age and sex. Results represent independent experiments. n= 5-10 mice per genotype per experiment unless otherwise indicated. Data are presented as violin plots representing kernel density distributions, with the interquartile range indicated. Scale bars: 50μm.

Next, we performed bulk RNA-seq on flow-sorted lung macrophages from *Tet2*-CHIP and control mice 14 days after bleomycin injury, to determine whether the heightened inflammatory milieu in CHIP lungs reflects altered macrophage activation. *Tet2*-CHIP macrophages exhibited broad transcriptional reprogramming, with induction of innate immune activation genes (*Casp4, Ifnb1, Tlr3, S100a8, Ccl11*), effector cytokines (*Csf2-Gm-csf, Ccl24, Osmr*), and inflammatory metabolic mediators (*Glrx, Abca1, Gpr65*). Concurrently, they upregulated profibrotic and matrix-remodeling programs, including MMPs (*Mmp12, Mmp13, Mmp3, Mmp27*), collagen-regulating factors (*Timp1, Sned1, Thbs1*), and growth modulators (*Ccn3, Gdf3, Gdf10, Fgf2, Wnt11, Sfrp1, Grem2*) **(Extended Data Fig. 4A, Supplementary Table 1)**. Several macrophage-intrinsic programs were suppressed, including antioxidant defenses (*Gsta4, Msrb3*), autophagy and vesicle-trafficking machinery (*Atg9a, Vamp7, Rab36*), and regulators of immune restraint (*Pik3ip1, Tnfsf18*), changes predicted to impair efferocytosis, metabolic homeostasis, and negative feedback control **(Extended Data Fig. 4A, Supplementary Table 1)**. Enrichment analysis supported these findings **(Extended Data Fig. 4B, Supplementary Table 2)**, revealing a persistently activated, poorly resolving macrophage state indicative of a maladaptive program aligned with the enhanced fibrosis observed in *Tet2*-CHIP lungs.

To further resolve macrophage states, we applied non-negative matrix factorization (NMF) to the bulk RNA-seq dataset, identifying two dominant metaprograms: NMF4, enriched in control macrophages, and NMF5, enriched in *Tet2*-CHIP macrophages **(Fig. 4B**, **Extended Data Fig. 4C, Supplementary Table 3)**. NMF4 was associated with homeostatic and reparative genes (*C1qa, Fcer1g, Lgals1*) and pathways related to immune surveillance and tissue maintenance **(Fig. 4B,C)**. In contrast, NMF5 was marked by inflammatory and stress-response genes (*Il1a, Ereg, Tlr2, Cybb, Tnfaip2, Tnfaip3*) and pathways supporting apoptotic regulation, metabolic adaptation, and inflammatory signaling **(Fig. 4B,C)**. Both clusters shared a common injury-response module (*Il1b, Cxcl1/2, Tnf, Spp1*), but *Tet2*-CHIP macrophages demonstrated a shift toward an amplified inflammatory and damage-responsive state.

To connect these transcriptional programs with specific macrophage populations, we leveraged scRNA-seq data from bleomycin-injured lungs^35^, which annotated an injury-responsive Spp1^+^ alveolar macrophage (AM) population that expands upon injury (**Extended Data Fig. 4D-F, Supplementary Table 4**). Using this dataset as a reference, deconvolution of *Tet2*-CHIP bulk RNA-seq revealed a significant enrichment of the Spp1^+^ AM signature **(Fig. 4D)** in *Tet2*-CHIP mice, indicating expansion of this injury-associated population. Immunofluorescence confirmed increased CD68^+^ Spp1^+^ macrophages in both *Tet2*- and *Asxl1*-CHIP mice compared with controls (**Fig. 4E,F**), the majority originating from mutant (CD45.2^+^) cells **(Fig. 4F)**. *Tet2*-CHIP lungs further showed an increase in IL-1β^+^ mutant macrophages (**Fig. 4G,H**), while both CHIP models exhibited expansion of Tgf-β1^+^ and Areg^+^ mutant macrophages **(Fig. 4I,J)**, linking CHIP-driven transcriptional reprogramming to a hyperinflammatory, profibrotic macrophage phenotype.

To evaluate whether analogous macrophage alterations occur in IPF patients with CHIP, we analyzed a publicly available, Universal 5’ Gene Expression scRNA-seq dataset from individuals with IPF that includes immune, epithelial, mesenchymal and endothelial cells **(Extended Data Fig. 4G-I)** ^36^. Using SComatic^37^, we predicted CHIP mutations and identified a subset of cases harboring mutations in the monocyte-macrophage lineage (6 cases predicted to carry CHIP mutations out of 27 total IPF cases) **(Extended Data Fig. 4G,H, Supplementary Table 5)**. Within this compartment, we identified an MDM-SPP1 population whose transcriptional profile closely matched the Spp1^+^ AMs identified in mice **(Fig. 4K, Supplementary Table 6)**. MILO analysis revealed significant enrichment of this MDM-SPP1 population in IPF patients with predicted CHIP mutations **(Fig. 4L)**. Differential expression analyses showed that CHIP-associated MDM-SPP1 cells upregulated pro-inflammatory and stress-response genes (*IL1B, TNFAIP3, S100A9/12, JUN, FOS, CEBPB*) as well as growth factors (*VEGFA, HBEGF, TGFB1, AREG*) **(Fig. 4M, Supplementary Table 7)**(hereafter *CHIP-signature*) and were enriched for NF-κB, MAPK, PI3K-Akt, myeloid activation, and Th17 differentiation pathways **(Fig. 4N)**. In contrast, MDM-SPP1 cells from non-CHIP IPF patients expressed higher levels of homeostatic and phagocytic genes (*CD36, C1QA/B/C, APOE, MT-ND3/4L/6*) **(Fig. 4M)** and were enriched for phagosome, lysosome, ER processing, and translation pathways **(Fig. 4N)**. Both groups shared wound-response and cytokine-signaling programs **(Fig. 4N)**. Notably, the CHIP-IPF upregulated gene signature localized almost exclusively to MDM-SPP1 macrophages in the annotated UMAP **(Fig. 4O)**, highlighting convergence on an analogous SPP1^+^ macrophage program across mouse and human CHIP.

To examine whether this CHIP-Signature (differentially upregulated genes in CHIP-associated MDM-SPP1 cells) could predict clinical outcomes we performed Cox proportional hazards analyses adjustment for age, sex, and batch effects. The *CHIP-Signature* genes were projected onto a publicly available dataset from broncho alveolar lavage (BAL)^38^. Patients were stratified into high- and low-signature groups based on aggregated expression scores. Importantly, individuals with high *CHIP-Signature* expression exhibited significantly worse overall survival (Median OS: 1176 days vs 574 days; p=3.11×10^−4^)(Fig. 4P) and a markedly increased hazard of death (HR= 2.70, 95% CI: 1.67-4.55).

Together, these findings demonstrate that CHIP-associated mutations amplify inflammatory and profibrotic macrophage programs, expand an injury-responsive SPP1^+^ population across mouse and human lung injury, and generate a transcriptional signature whose expression associates with worse clinical outcomes in independent IPF cohorts, underscoring the pathogenic and prognostic significance of CHIP-driven macrophage reprogramming.

### CHIP promotes fibroblast activation and epithelial dysfunction in pulmonary fibrosis

To determine whether the accumulation of CHIP-associated Spp1^+^ macrophages with pro-inflammatory and profibrotic features translates into altered parenchymal remodeling, we next examined fibroblast and alveolar epithelial responses in *Tet2*- and *Asxl1*-CHIP mice after injury. Immunofluorescence analysis of lung sections from *Tet2*- and *Asxl1*-CHIP mice revealed coordinated alterations in profibrotic fibroblast populations 14 days post bleomycin injury. Both models exhibited a significant increase in the frequency of αSMA^+^ fibroblasts, with a larger subset co-expressing Runx1 in CHIP-mice, indicating expansion of activated myofibroblasts **(Fig. 5A,B)**. The injury-associated Cthrc1^+^ fibroblast population was increased in *Tet2*-CHIP lungs but not in *Asxl1*-CHIP mice, whereas the fraction of Cthrc1^+^Runx1^+^ fibroblasts was significantly elevated in both models **(Fig. 5C,D)**. By contrast, intermediate Sfrp1^+^ fibroblast frequencies were largely preserved, with a modest decrease observed in the *Asxl1* model **(Fig. 5E,F)**. To determine whether CHIP macrophages are sufficient to induce these fibroblast activation changes, we co-cultured primary wild-type lung fibroblasts with macrophages isolated from *Tet2*- or *Asxl1*-CHIP mice. Consistent with our *in vivo* findings, CHIP macrophages significantly increased the frequency of αSMA^+^ fibroblasts, expanded αSMA^+^Runx1^+^ fibroblasts and promoted the emergence of Cthrc1^+^ fibroblasts relative to control macrophages **(Fig. 5G-J)**. Parenchymal epithelial responses were also altered *in vivo*. In *Tet2*-CHIP lungs, but not *Asxl1*, the frequency of Spc^+^ AT2 cells was reduced, whereas both models exhibited an accumulation of Krt8^+^ intermediate cells, including Krt8^+^Aqp5^+^ transitional states, without a significant change in mature Aqp5^+^ AT1 cell abundance **(Fig. 5K-O)**. Proliferation analysis demonstrated an increased frequency of cycling Ki67^+^Krt8^+^ cells in both models, with no corresponding increase in Ki67^+^ Spc^+^ AT2 cells **(Fig. 5M,P and Extended Data Fig. 5A,B)**, suggesting a skewing toward transitional, injury-associated epithelial states. In organoid cultures, AT2 cells co-cultured with mutant macrophages formed significantly more and larger organoids than those cultured with wild-type macrophages, and these organoids accumulated an expanded Krt8^+^ transitional population **(Fig. 5Q-S)**. Altogether, these data demonstrate that CHIP, and CHIP-associated macrophages in particular, are sufficient to directly reprogram fibroblast and epithelial behaviors toward the maladaptive states observed in CHIP fibrotic lungs.

**Figure 5:**
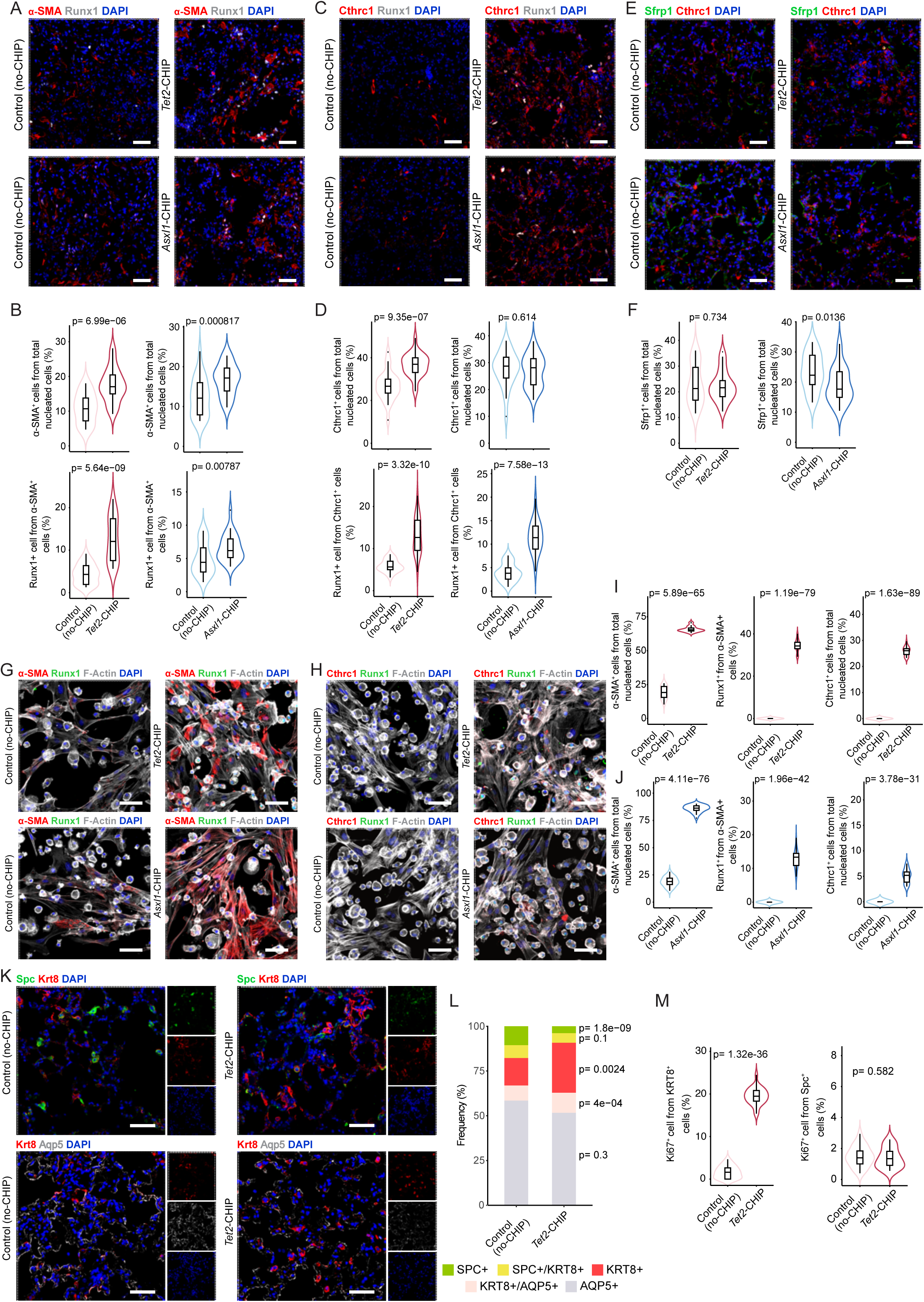
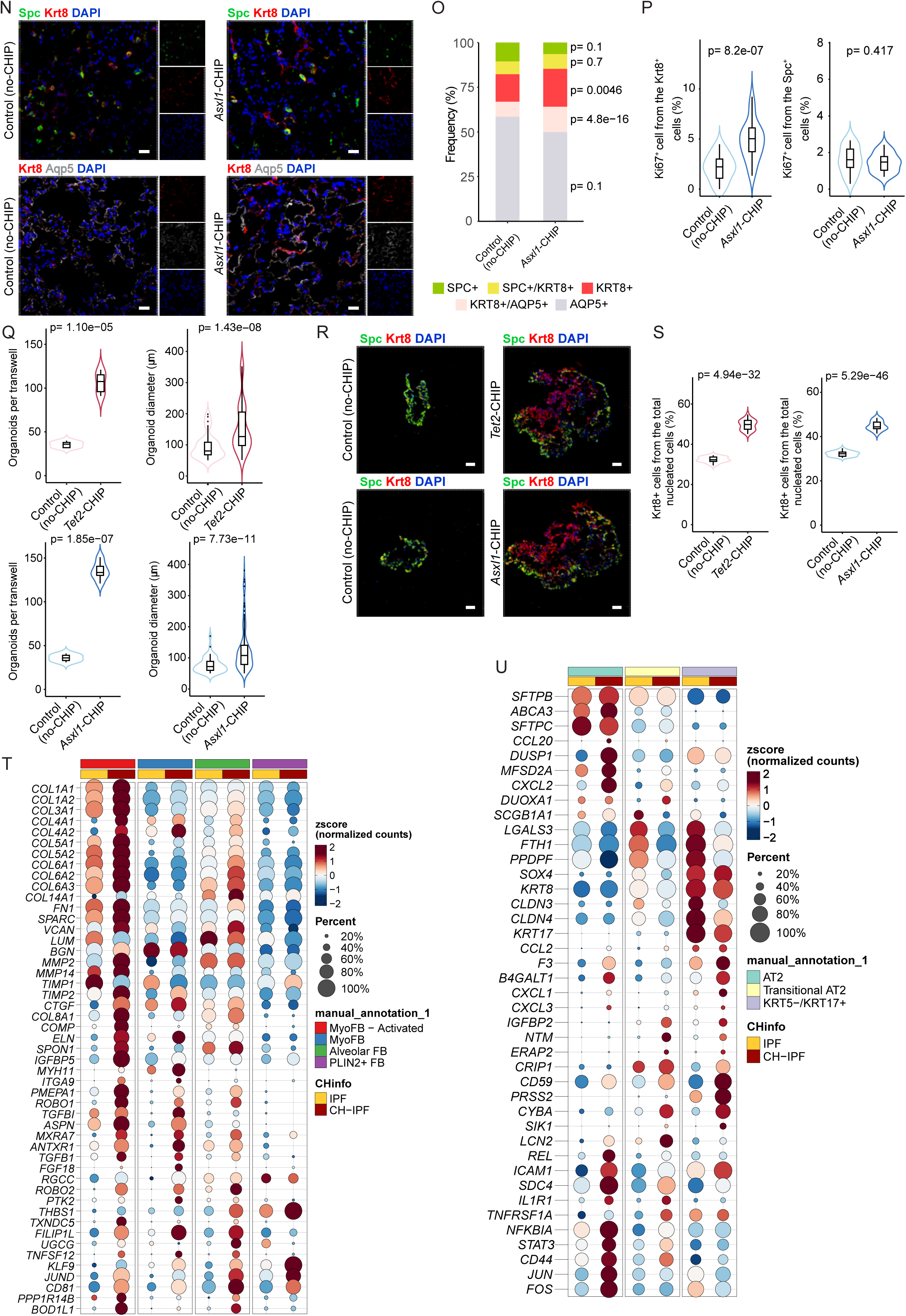
CHIP promotes fibroblast activation and epithelial dysfunction in pulmonary fibrosis. **a-f,** Representative IF images and their statistical quantification of lung sections from Control-no-CHIP, *Tet2-* and Asxl1-CHIP mice showing alpha-Smooth Muscle Actin (a-SMA), Runx1, Cthrc1, and Sfrp1. **g,h** Representative IF images from *in vitro* co-culture experiments of mouse wild-type fibroblasts and macrophages from Control-no-CHIP, *Tet2-* and Asxl1-CHIP mice, stained for a-SMA or Cthrc1, Runx1, and F-actin (Phalloidin). **i,j,** Quantification of the IF images shown in **g,h**. **k.** Representative IF images showing staining for Surfactant Protein C (Spc), Keratin 8 (Krt8) and Aquaporin 5 (AQP5) in lungs from Control-no-CHIP and *Tet2-*CHIP mice. **l.** Quantification of the IF images shown in **k**. **m,** Quantification of proliferating Spc^+^ cells and Krt8^+^ cells assessed by Ki67 expression from Control-no-CHIP and *Tet2-*CHIP **n.** Representative IF images showing staining for Spc, Krt8 and Aqp5 in lungs from Control-no-CHIP and *Asxl1-*CHIP mice. **o.** Quantification of the IF images shown in **n**. **p,** Quantification of proliferating Spc^+^ cells and Krt8^+^ cells assessed by Ki67 expression from Control-no-CHIP and *Asxl1-*CHIP. **q-s,** Organoid co-culture of Spc^+^ AT2 cells with macrophages from Control-no-CHIP, *Tet2-* and Asxl1-CHIP mice lung tissue in the presence of stromal support. **q,** Quantification of colony formation efficiency and size of organoid 14 days post plating; **r,** Representative IF images of the organoid co-culture stained for Spc and Krt8 14 days post plating. **s,** Quantification of the accumulation of Krt8+ intermediate epithelial cells in AT2 organoids coculture. **t,** Dot plot depicting relative expression of ECM structural components, collagen-processing enzymes, drivers of fibroblast activation, mediators of matrix assembly and cytoskeletal contractility in fibroblasts populations from IPF patients with or without predicted CHIP mutations. **u,** Dot plot depicting relative expression of inflammatory mediators, injury-responsive genes and structural components in alveolar epithelial cell populations from IPF patients with or without predicted CHIP mutations. Results represent independent experiments. n= 5-10 mice per genotype per experiment unless otherwise indicated. Data are presented as violin plots representing kernel density distributions, with the interquartile range indicated. Scale bars: 50μm.

Given the conserved macrophage phenotype in mouse and human CHIP, we next asked whether downstream fibroblast and epithelial remodeling programs were likewise altered in IPF patients carrying CHIP **(Fig. 5T,U**; **Extended Data Fig. 4G and Extended Data Fig. 5C,D)**. Across fibroblast states, CHIP was associated with pan-mesenchymal reprogramming toward a more activated, fibrogenic state consistent with the fibroblast phenotypes observed in our mouse models **(Fig. 5A-F)**. In myofibroblast-enriched clusters, CHIP-associated cells showed increased relative expression of multiple ECM structural components, collagen-processing enzymes (*MMP14, MMP2, TIMP2, TXNDC5*), key drivers of fibroblast activation, (*TGFB1, CTGF, PMEPA1, and ROBO1/ROBO2*) as well as mediators of matrix assembly (*ELN, FN1, SPON1, CTGF*) and cytoskeletal contractility (*MYH11, PTK2, FILIP1L*) **(Fig. 5T)**. GSVA analyses were concordant with these transcriptional shifts **(Extended Data Fig. 5E,F, Supplementary Table 8,9)**. Alveolar fibroblasts and Plin2^+^ fibroblasts, which support alveolar maintenance and early repair, exhibited modest but coordinated increases in extracellular matrix and matricellular components (*VCAN, SPON1, THBS1* and several collagen isoforms) together with elevated expression of regulatory and stress-adaptive transcriptional programs (*KLF9, JUND, RGCC*), consistent with a transition from a homeostatic to a more injury-primed reactive state **(Fig. 5T)**. GSVA pathway enrichment mirrored these transcriptional shifts **(Extended Data Fig. 5G, Supplementary Table 10)**.

Across epithelial subpopulations, CHIP-associated cells displayed transcriptional signatures consistent with heightened inflammatory sensitivity, impaired differentiation, and stress-adaptive remodeling, paralleling the maladaptive epithelial phenotypes observed in the mouse models. In AT2 cells, relative increases in inflammatory mediators and stress-responsive genes (*ABCA3, DUSP1, ICAM1, IL1R1, TNFRSF1A, STAT3, JUN/FOS, LCN2*) were accompanied by GSVA enrichment for inflammatory activation, NF-κB/MAPK signaling, and perturbed epithelial differentiation and survival programs **(Fig. 5U and Extended Data Fig. 5H, Supplementary Table 11)**. Transitional AT2 cells similarly showed elevated expression of injury-responsive mediators (*IGFBP2, LCN2, IL1R1, STAT3, CD44*) with concomitant reduction of homeostatic genes (*SCGB1A1, FTH1*), consistent with pathway enrichment for MAPK/JNK activation, Notch dysregulation, epithelial apoptosis, and DNA damage responses, features characteristic of stalled or maladaptive differentiation **(Fig. 5U and Extended Data Fig. 5I, Supplementary Table 12)**. Krt5^−^/Krt17^+^ aberrant basaloid cells demonstrated increased expression of chemokines and injury-response genes (*CXCL1, ICAM1, F3, CD59*) together with reduced junctional and structural markers (*LGALS3, CLDN3/4*), consistent with enrichment of interferon- and chemokine-driven signaling and cytoskeletal remodeling pathways **(Fig. 5U and Extended Data Fig. 5J, Supplementary Table 13)**. Collectively, these results show that CHIP reprograms fibroblast and epithelial responses toward fibrosis-prone states in both mice and humans, further reinforcing its role as a modifier of parenchymal repair and a potential contributor to fibrotic disease progression.

### CHIP establishes a primed parenchymal environment in the absence of overt injury

We next evaluated whether CHIP is sufficient to induce a subclinical alteration in lung parenchymal organization and cellular composition even in the absence of injury.

Histological evaluation of the *Tet2* and *Asxl1*-CHIP mouse models (hematoxylin–eosin staining and Masson’s trichrome) revealed no overt tissue damage or altered collagen accumulation compared to control mice **(Fig. 6A,B and Extended Data Fig. 6A)**. However, CHIP lungs displayed focal morphological abnormalities, including sporadic enlarged cells resembling lipid-laden macrophages in the alveolar space **(Fig. 6A,B)**. In addition, both *Tet2*- and *Asxl1*-CHIP mice demonstrated a modest but statistically significant increase in alveolar wall thickness compared with their respective controls **(Fig. 6C)**, indicating early perturbations in parenchymal integrity before fibrotic challenge. Immunofluorescence analysis further revealed modest but significant increases in Col1a1^+^ matrix deposition in *Tet2*- and *Asxl1*-CHIP lungs compared with their respective controls, accompanied by a positive correlation between Col1a1 coverage and areas enriched in macrophages **(Fig. 6D-F)**. In parallel, epithelial staining demonstrated an elevated frequency of transitional (Krt8^+^ and Spc^+^/Krt8^+^) epithelial cells in CHIP-mice, together with increased Ki67 expression within these intermediate populations, **(Fig. 6G-I)** whereas AT2 cell frequency and proliferation remained unchanged **(Fig. 6H and Extended Data Fig. 6B)**. Consistent with our observations in injured lungs, CHIP-mice also exhibited a baseline expansion of Spp1^+^ macrophages, the majority derived from the mutant (CD45.2^+^) compartment **(Fig. 6J-L)**. Notably, the abundance of Krt8^+^ transitional epithelial cells correlated with areas of Spp1^+^ macrophages abundance **(Fig. 6M,N)**, suggesting that even in the absence of overt injury, CHIP promotes a microenvironment in which Spp1^+^ macrophages accumulate in areas of active epithelial regeneration with presence of epithelial intermediate cells. In this context, transcriptional profiling of lung macrophages in *Tet2*-CHIP mice further revealed broad activation of inflammatory and stress-responsive regulatory programs. Key macrophage transcription factors, including C/EBP family members, STAT3, NF-κB components, AP-1 factors, IRFs, and SMAD-dependent mediators, displayed increased inferred activity **(Fig. 6O)**. Similarly, enrichment analysis revealed activation of transcriptional programs consistent with a primed, proinflammatory state in *Tet2*-CHIP macrophages **(Extended Data Fig. 6C, Supplementary Table 14)**.

**Figure 6:**
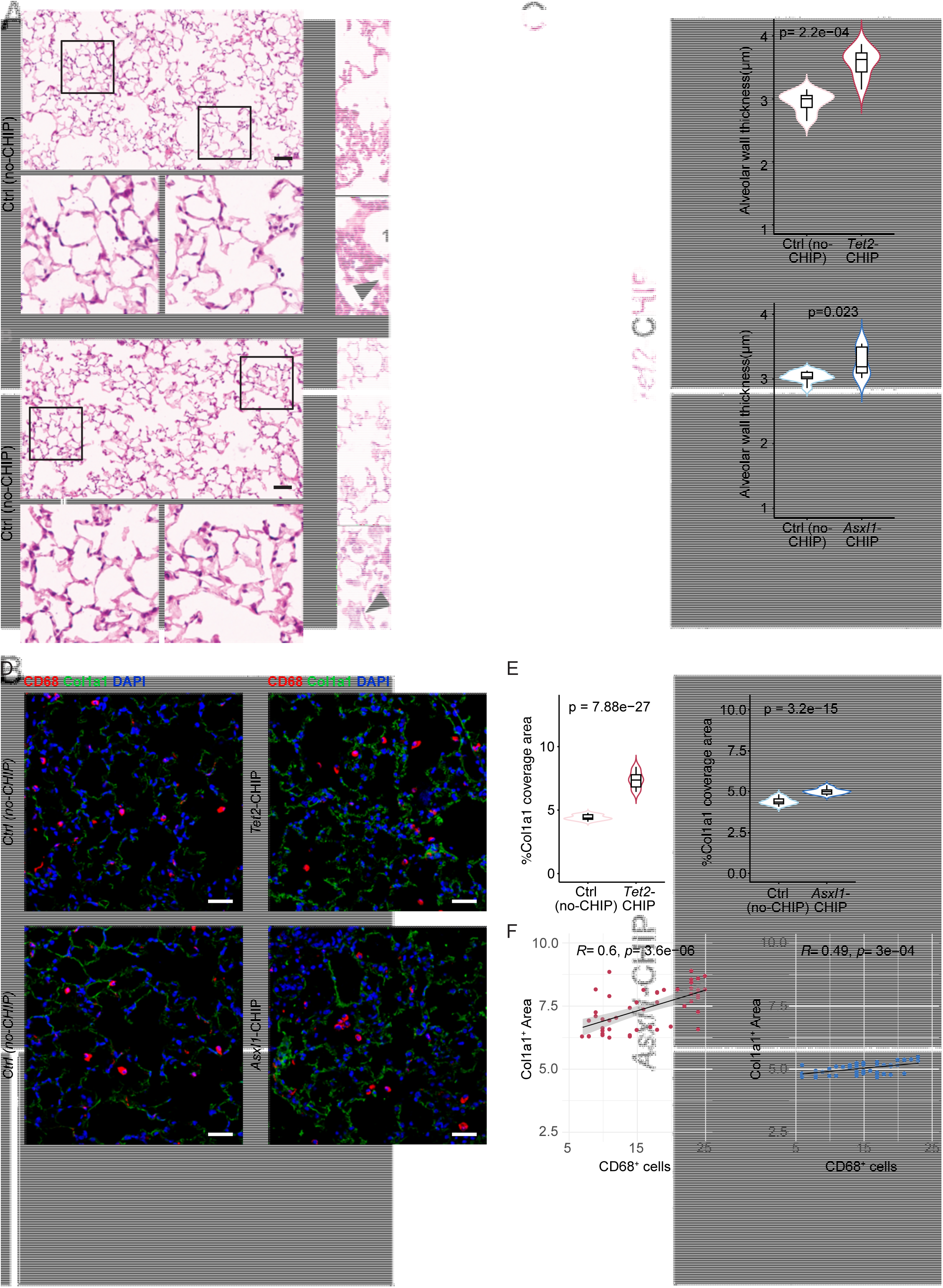
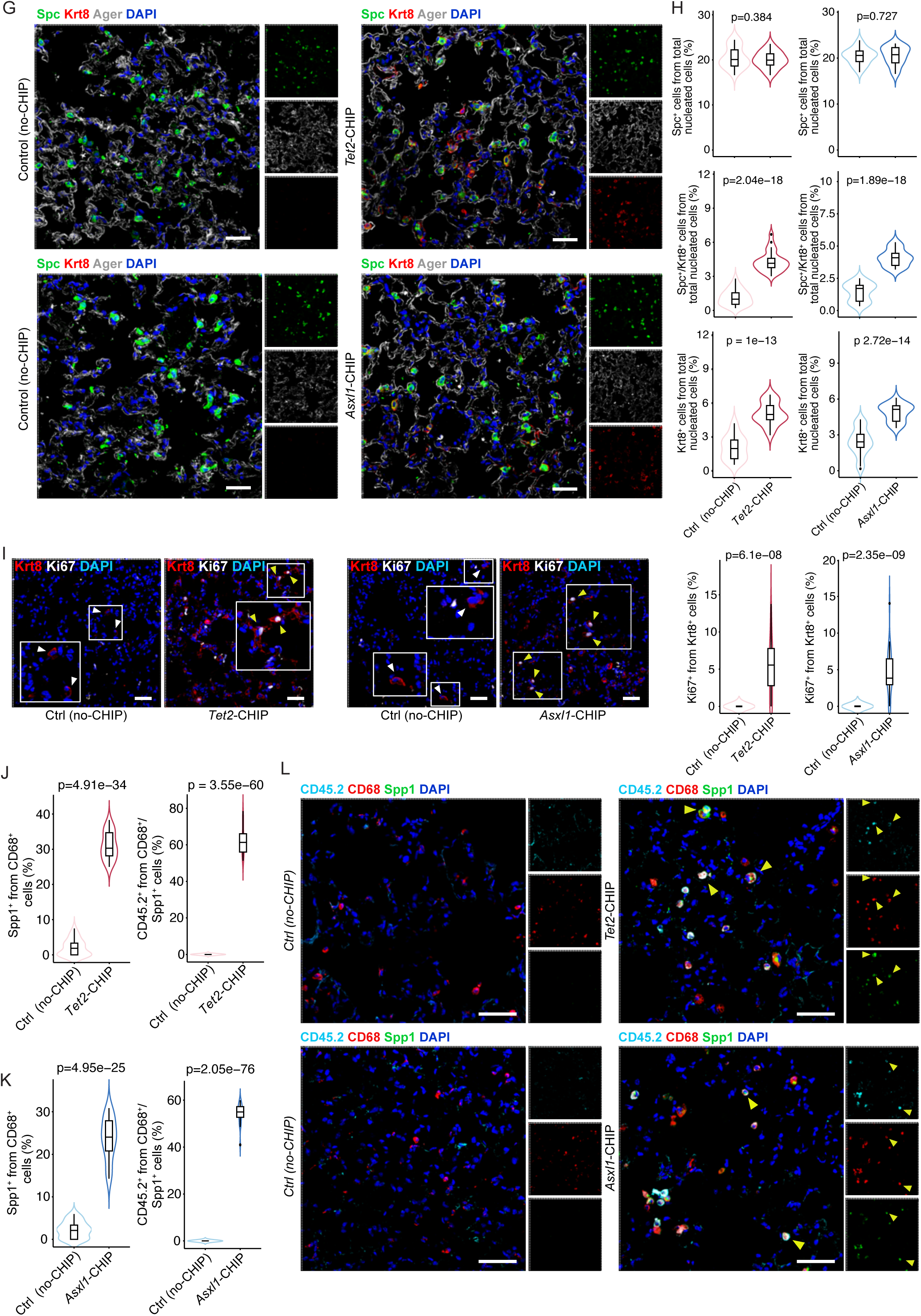
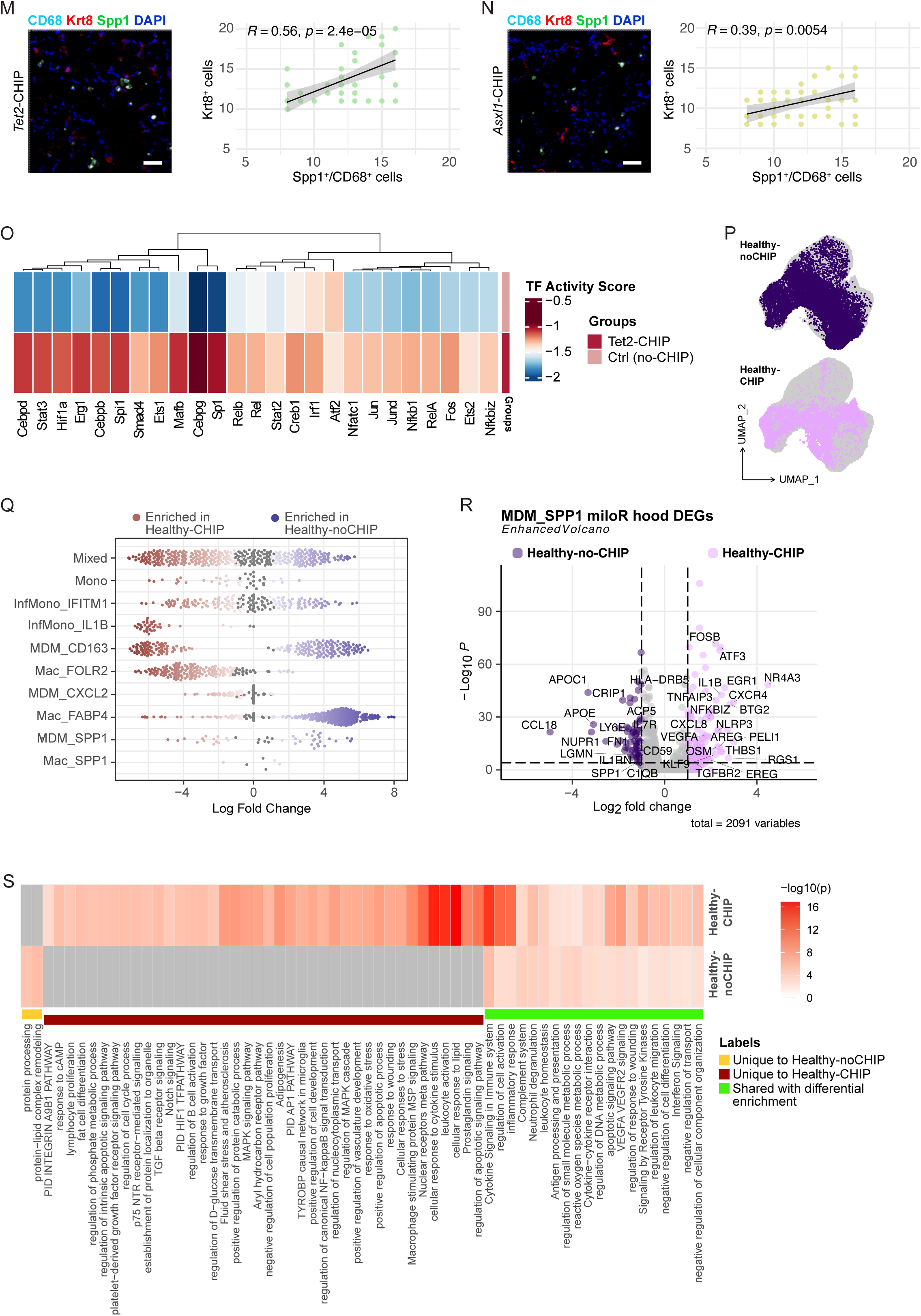
CHIP establishes a primed parenchymal environment in the absence of overt injury. **a,b,** Representative images of lung histology stained with Hematoxylin and Eosin (H&E) in lungs from *Tet2-*, *Asxl1-*CHIP and control noCHIP mice in the absence of injury. Enlarged representative H&E images represent detailed visualization of alveolar architecture. Arrows highlight large cells observed in *Tet2-* and *Asxl1-*CHIP mice. **c,** Quantification of alveolar wall thickness based on H&E staining in uninjured *Tet2-*, *Asxl1-*CHIP and control noCHIP lungs. **d,** Representative IF images showing CD68^+^ cells and Col1a1 coverage area in Control-no-CHIP, *Tet2-* and *Asxl1-*CHIP in the absence of injury. **e,f,** Statistical quantification of Col1a1 coverage area (**e**) and Pearson correlation analysis (**f**) between Col1a1 coverage area and CD68^+^ cell counts, where each dot represents a field (HPF). **g,h** Representative IF images (**g**) and quantification (**h**) of alveolar epithelial cells stained for Spc, Krt8, and Advanced Glycation End-products Specific Receptor (AGER) from Control-no-CHIP, *Tet2-* and *Asxl1-*CHIP in the absence of injury. **i,** Statistical quantification and representative IF images of Krt8^+^ cell proliferation assayed by Ki67 co-localization in the lungs from Control-no-CHIP, *Tet2-* and *Asxl1-*CHIP in the absence of injury. White arrowheads point to non-proliferating Krt8+ cells, while yellow arrowheads point to double positive Ki67^+^ Krt8^+^ cells. **j-l,** Statistical quantification of the co-localization of CD68^+^ and CD45.2^+^ macrophages with Spp1 in the lungs from Control-no-CHIP, *Tet2-* and *Asxl1-*CHIP in the absence of injury (**j,k**) and representative IF images (**l**). **m,n,** Pearson correlation analysis between CD68^+^/Spp1^+^ macrophages and Krt8^+^ cell counts, where each dot represents a field (HPF) from Control-no-CHIP, *Tet2-* and *Asxl1-*CHIP in the absence of injury. **o,** Transcription Factor (TF) activity analysis based on lung macrophage bulk RNA-seq data from Control-no-CHIP and *Tet2-*CHIP in the absence injury. **p,** UMAP plot showing CHIP status (Healthy-noCHIP or Healthy-CHIP), identified by single-nucleotide variant (SNV) calling, in the healthy human scRNA-seq dataset (GSE227136). **q,** Beeswarm plot of the log2(fold change) in the Milo neighborhoods between healthy individuals with and without predicted CHIP mutations, grouped by cell type. Neighborhoods showing a significant change in cellular abundance are colored. **r,** Volcano plot showing differentially expressed genes (DEGs) between human healthy individuals with or without predicted CHIP mutation within the MDM_SPP1 cluster, restricted to cells from Milo neighborhoods identified as significant in panel. Genes with P-value <0.01 and log2(fold change) > 1 are color-coded. **s,** Comparative functional enrichment analysis of the gene signatures identified in panel r between healthy individuals with and without predicted CHIP mutations. Results shown represent independent experiments. n= 4-6 mice per genotype per experiment unless otherwise indicated. Data are presented as violin plots representing kernel density distributions, with the interquartile range indicated. Scale bars: 50μm.

To assess whether CHIP influences macrophage states in the uninjured human lung, healthy control donors from the same scRNA-seq dataset previously described were analyzed **(Extended Data Fig 4. G,H)**. After calling SNVs and inferring CHIP status, we mapped CHIP-associated and non-CHIP macrophages onto the annotated myeloid UMAP **(Fig. 6P)**. MILO analysis identified several macrophage clusters with altered abundance patterns, though, unlike in IPF, the MDM-SPP1 population did not show CHIP-specific expansion **(Fig. 6Q)**. Nonetheless, differential expression within the MDM-SPP1 compartment showed that CHIP-associated cells upregulated a broad inflammatory and stress–response program, including immediate-early transcription factors (*EGR1, FOSB, ATF3, NR4A1/2/3*), cytokine/NF-κB pathway mediators (*IL1B, TNFAIP3, NFKBIZ, CXCL2/8*), metabolic and survival regulators (*NAMPT, PFKFB3, PDK4, MCL1*), and growth-factor/AREG–HBEGF–VEGFA axis components; while downregulated genes were enriched for homeostatic and immunoregulatory functions (*APOE, APOC1, C1QB/C, IL1RN, CTSL, PLA2G7*) **(Fig. 6R, Supplementary Table 15)**. These transcriptional shifts were supported by pathway enrichment analysis, which demonstrated that CHIP macrophages preferentially engaged AP-1/NF-κB/MAPK signaling, HIF-1 and oxidative-stress programs, TGF-β and Notch-associated pathways, and processes related to cytokine responsiveness, lipid handling, apoptotic signaling, and leukocyte activation, whereas pathways shared across CHIP and non-CHIP macrophages reflected core macrophage biology such as antigen presentation, interferon signaling, complement activity, and neutrophil degranulation **(Fig. 6S)**. Together, these data indicate that even in the absence of overt disease, CHIP is associated with a transcriptionally primed, pro-inflammatory and stress-responsive macrophage state in the human and mouse lung, establishing a primed immune landscape that may lower the threshold for maladaptive repair once injury occurs.

## Discussion

Clonal hematopoiesis of indeterminate potential (CHIP) has emerged as an important determinant of systemic organ health, with established links to aging-associated diseases, and mortality^9,14,16^. These observations have reframed CHIP from a benign age-related phenomenon to an active modifier of inflammatory disease biology. However, its role in idiopathic pulmonary fibrosis (IPF) has remained largely unexplored, despite IPF being a prototypical age-associated disorder characterized by chronic tissue injury, immune dysregulation, and poor clinical outcomes.

Here, we show that CHIP represents a clinically and biologically relevant modifier of IPF. In contrast to the general aging population, IPF was associated with a distinct distribution of CHIP mutations, characterized by relative enrichment of non-*DNMT3A* variants, including *ASXL1* and *PPM1D*, a mutation-pattern consistent across age, sex, and smoking strata, supporting a model in which IPF-associated inflammation modifies clonal selection beyond known demographic risk factors. Moreover, CHIP driven by non-*DNMT3A* mutations, and particularly by larger clones, was associated with higher odds of IPF, indicating the presence of a selective CHIP endotype with distinct biological and clinical implications.

Mechanistically, our data support a conserved immune-driven pathway linking CHIP to fibrotic lung remodeling. In murine models, CHIP exacerbated bleomycin-induced pulmonary fibrosis was accompanied by increased inflammatory signaling and parenchymal damage. CHIP was associated with macrophage reprogramming toward an inflammatory, profibrotic state, including expansion of an injury-responsive SPP1^+^ population. This macrophage state was conserved across species, sufficient to alter fibroblast and epithelial behavior *ex vivo*, and characterized by a transcriptional signature associated with poorer outcomes in independent IPF cohorts, positioning CHIP-associated macrophages as a central node linking hematopoietic mutations to immunofibrotic remodeling.

Downstream of immune reprogramming, we observed coordinated maladaptive responses in fibroblast and epithelial compartments. CHIP was associated with enhanced fibroblast activation, matrix-remodeling programs, and altered epithelial differentiation trajectories, including expansion of transitional epithelial states that are implicated in the fibrotic process contributing to fibroblast activation^39^. Notably, aspects of this parenchymal reprogramming were detectable even in the absence of overt lung injury, indicating that CHIP can establish a primed, inflammation-biased tissue environment that lowers the threshold for maladaptive repair.

Rather than reflecting passive age-related clonal drift alone, our data support a model in which systemic clonal composition is shaped by chronic disease-associated selective pressures. In this framework, the repetitive injury to the lung acts as a source of sustained inflammatory and stress signals that may favor expansion of specific mutant hematopoietic clones in the bone marrow. In turn, CHIP-derived immune cells recruited to the lung amplify inflammatory and profibrotic responses, exacerbating tissue injury. This bidirectional interaction provides a coherent explanation for the circulating enrichment of particular CHIP mutations in IPF and their disproportionate impact on lung pathology. This may explain the recent reported increase in clonal hematopoiesis in DDR (DNA Damage Repair) genes, including *PPM1D*, recently observed in patients undergoing lung transplant for a myriad of chronic lung diseases including ILDs^40^.

These findings have several clinical implications. First, we identify CHIP as a potential mechanistic biomarker of an inflammatory-driven IPF endotype, which may help explain heterogeneity in therapeutic responses observed across prior clinical trials. Immunomodulatory strategies that failed in unselected IPF populations, such as tumor necrosis factor inhibition in the Etanercept trial (NCT00063869)^41^, may warrant re-evaluation in CHIP-stratified cohorts, particularly considering evidence from the CANTOS trial showing selective benefit of inflammatory pathway modulation in CHIP carriers^42^. Conversely, the adverse outcomes observed in the PANTHER-IPF trial^43^ with thiopurine-based immunosuppression, agents known to induce DNA damage, may reflect unrecognized interactions with clonal hematopoiesis that favor expansion of pathogenic clones, providing a potential mechanistic explanation for their unexpected harm. Collectively, these observations suggest that prior negative trial results may reflect the absence of biologically informed patient stratification and that targeted anti-inflammatory approaches guided by CHIP status could benefit selected subsets of patients with IPF. Second, whereas CHIP is typically not considered a direct therapeutic target, in a rapidly progressive and fatal disease such as IPF, the potential to mitigate disease progression may outweigh the risks of intervention.

More broadly, given the established association between CHIP and cardiovascular disease, IPF patients with CHIP may benefit from intensified cardiovascular risk assessment and management, underscoring the need for coordinated, interdisciplinary care. CHIP may also influence progression from interstitial lung abnormalities to overt fibrosis, supporting closer longitudinal monitoring. Conversely, therapies that promote clonal selection within the hematopoietic system, including certain chemotherapeutics, may carry unintended pulmonary consequences in susceptible individuals. In addition, systematic evaluation of CHIP status in both lung transplant donors and recipients may be warranted, as CHIP-associated immune reprogramming could plausibly influence graft inflammation, chronic rejection, and long-term allograft function. At the population level, the high prevalence of CHIP in aging individuals and association with adverse outcomes across organ systems positions it as a candidate biomarker for risk-based screening and preventive strategies across pulmonary, cardiovascular, and oncologic domains.

Our study has several limitations. Although our data support a pathogenic association between CHIP-associated immune reprogramming and IPF, causal inference in humans remains constrained by the observational nature of clinical datasets and unaccounted for confounders. CHIP status in lung immune cells was inferred from 5′ scRNA-seq data, which has limited sensitivity for small-sized clones and likely underestimates CHIP prevalence^44^. In addition, experimental studies focused on selected CHIP mutations may not capture the full diversity observed in patients. Prospective studies will be required to determine whether CHIP-informed stratification improves therapeutic response, prognostication, or clinical decision-making in IPF and related fibrotic lung diseases.

In conclusion, our data extend the concept of CHIP beyond passive age-associated clonal drift. The distinct mutational patterns observed in IPF, together with experimental evidence of immune-parenchymal crosstalk, support a model in which chronic tissue injury and inflammation actively shape the hematopoietic mutational landscape with clinically meaningful consequences. In this framework, CHIP is both a product and a driver of disease biology, linking aging, immune dysregulation, and organ-specific pathology in fibrotic lung disease.

## Acknowledgments

We thank all current and former members of the Saez Laboratory, and the Pardo-Saganta Lab; the people involved in the maintenance of mouse colonies; the CIMA and ILH flow cytometry core facilities; the histology core at CIMA and aRhitmó. We gratefully acknowledge all colleagues that contributed technically and/or intellectually and the patients that kindly donated tissue samples that were key for the completion of this work. We are grateful to the late Tobias Welte for his vision and contribution to the project. We thank the Hannover Unified Biobank at MHH for processing and storage of the biological samples. This research was supported in part by the Instituto de Salud Carlos III ISCIII PI17/01346 and PI20/00152, co-funded by ERDF,“A way to make Europe” (to BSa); Gobierno de Navarra (0011-3638-2020-000011 and 0011-3597-2020-000005) co-funded by the ERDF through the Operative Program 2014-2020 of Navarra to BSa; an AECC Junior Investigator Grant exp. AIO16163636SAEZ (to BSa), a Ramon y Cajal Award RYC-2015-18580 co-founded by FSE (to AP-S), and a grant from Gobierno de España SAF2017-89908-R (MINECO/AEI/FEDER, UE.) (to AP-S). FHH was supported in part by grants of the German Research Council (DFG): HE6233/15-1 and 16-1, project number 517204983, the Leibniz Association (XPAND-HSC, TICKTOCK) and by the EPPERMED2025-134 (HOPE-Consortium). JCS was supported by the Else Kröner-Fresenius Foundation (2023_EKCS.18) and the German Center for Lung research (FKZ 82DZL002C1 & FKZ 82DZLT82C1). SBM was supported in part by the NIH/NHLBI (K23HL150331).

## Author contribution

**DL** designed and performed the experiments, analyzed and interpreted the data, and edited the manuscript; **ACV** and **PG-O**, designed and performed the experiments, analyzed and interpreted the data; **SBM, BSS, AP and BSe** provided human samples and data from the MGH and MHH cohorts, **IV-U, PA, BA, MN and LC** performed the genomic sequencing, analyzed and interpreted the data; **AT and NB** optimized and performed *in vitro* co-culture experiments and helped with the analysis; **PSV, LV and EP** performed *in vivo* experiments and analysis, and helped with the interpretation of the data; **AV and PSM** performed RNA sequencing, analyzed and interpreted the data; **MJC and, FP** contributed to the sequencing of human samples from the MGH cohort and the interpretation of the data; **PGM** contributed vital new reagents and or analytical tools and participated in the interpretation of the data; **WS** contributed in the interpretation of the data and edited the manuscript: **JCS** and **FHH** conceived the study, provided data derived from the sequencing of IPF patients from the MHH cohort, interpreted the data, provided feedback on the analysis of human data and edited the manuscript; **BSa and AP-S** conceived the study, designed and performed the experiments, analyzed and interpreted the data, supervised the project, coordinated the entire team, and wrote the manuscript.

---

**Extended data Fig. 1:**
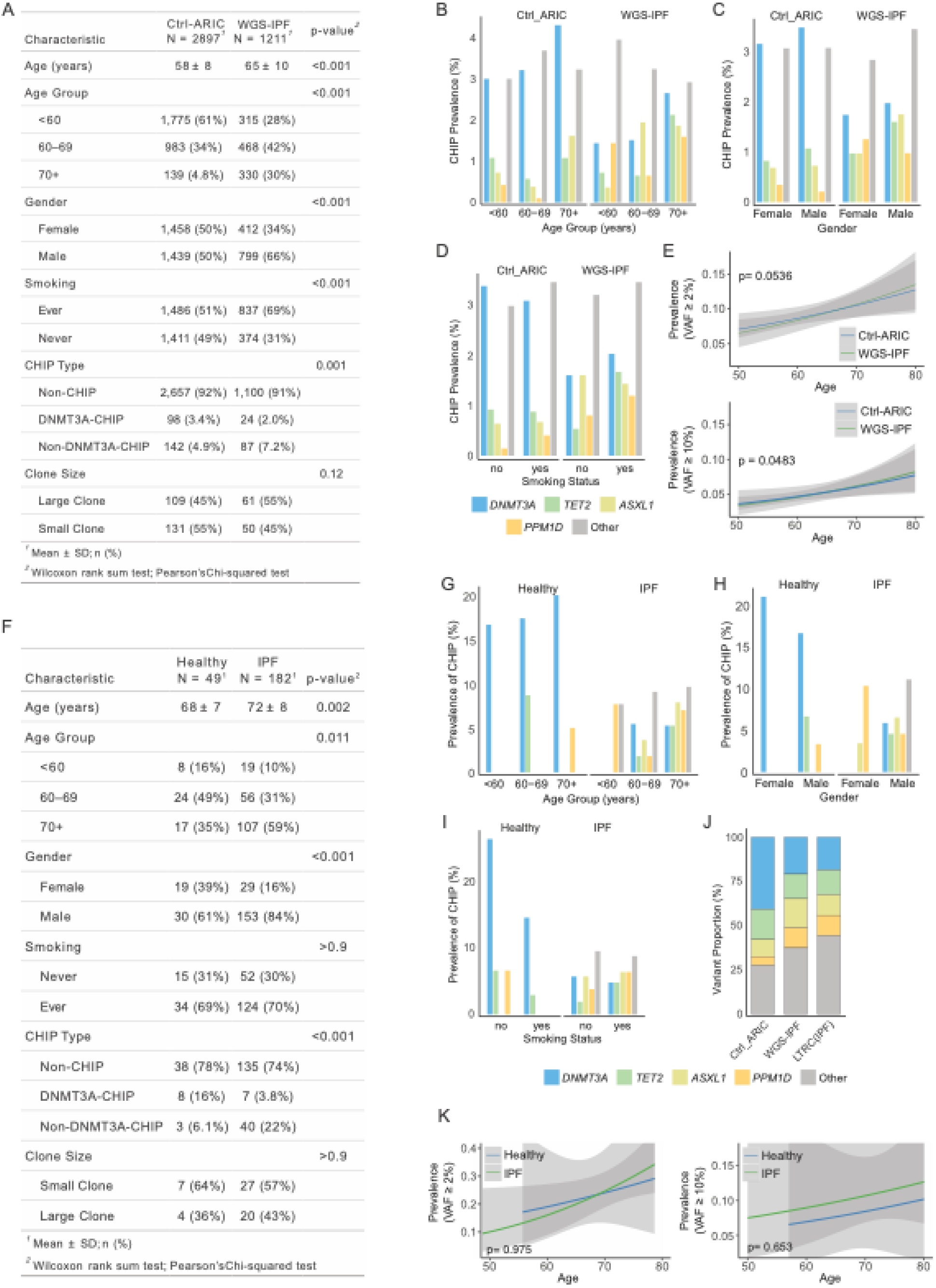
Contingency tables and demographic distribution of CHIP variants in TOPMed and validation cohorts. **a,** Contingency table summarizing demographic characteristics of the TOPMed WGS ARIC and PFWGS cohorts. **b-d,** Distribution of CHIP variants by **b,** age; **c,** sex; and **d,** smoking status in the TOPMed WGS ARIC and PFWGS cohorts. **e,** Age-associated clonal expansion, measured by variant allele fraction (VAF), in the TOPMed WGS ARIC and PFWGS cohorts. **f,** Contingency table summarizing demographic and clinical characteristics of the validation cohort. **g-i,** Distribution of CHIP variants by **g,** age; **h,** sex; and **i,** smoking status in the validation cohort. **j**, Relative abundance of CHIP-associated gene variants among ARIC, PFWGS and LTRC (IPF cases, n=248) cohorts **k,** Age-associated clonal expansion, measured by VAF, in the validation cohort.

**Extended data Fig. 2:**
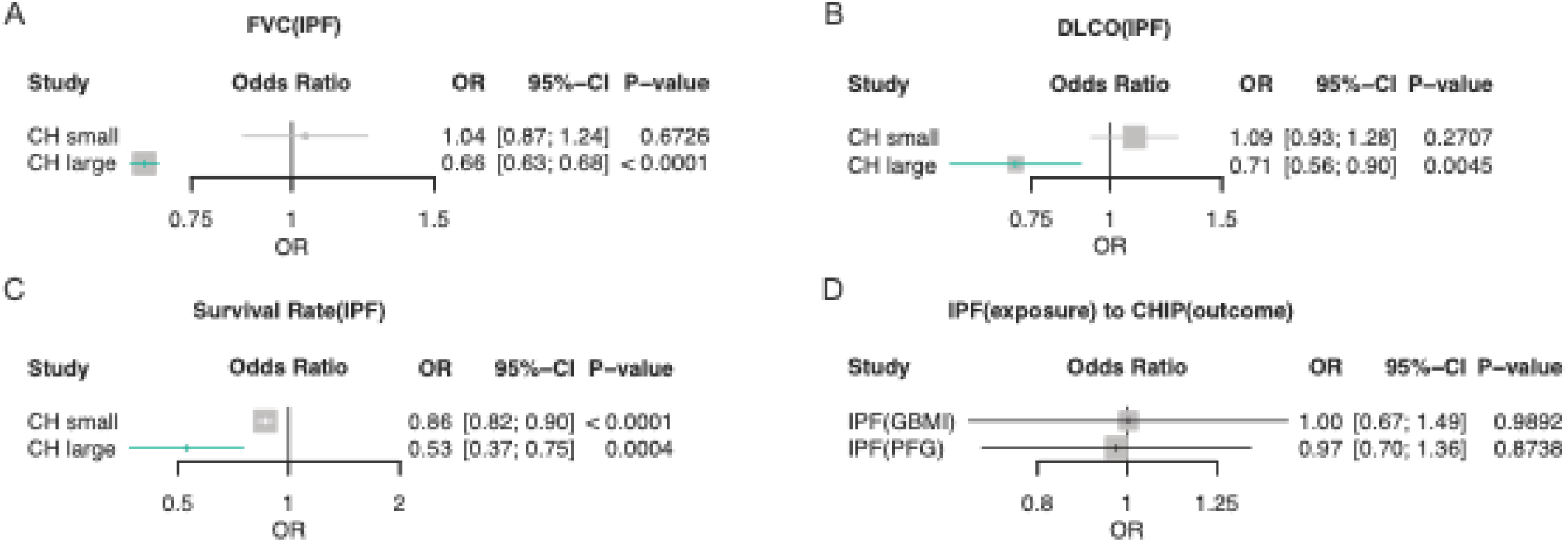
Mendelian randomization estimates for the effect of CHIP clone size on IPF clinical outcomes. **a-c,** MR exploring the association between CHIP clone size and clinical outcomes within the PFG cohort: (**a,** Forced Vital Capacity (FVC), **b,** Diffusing Capacity of the Lung for Carbon Monoxide (DLCO), and **c,** Transplant-free Survival Rate.). **d,** Reverse MR analysis assessing the association between IPF and CHIP risk. Results are presented as odds ratios (ORs) with 95% CIs and p-values.

**Extended data Fig. 3:**
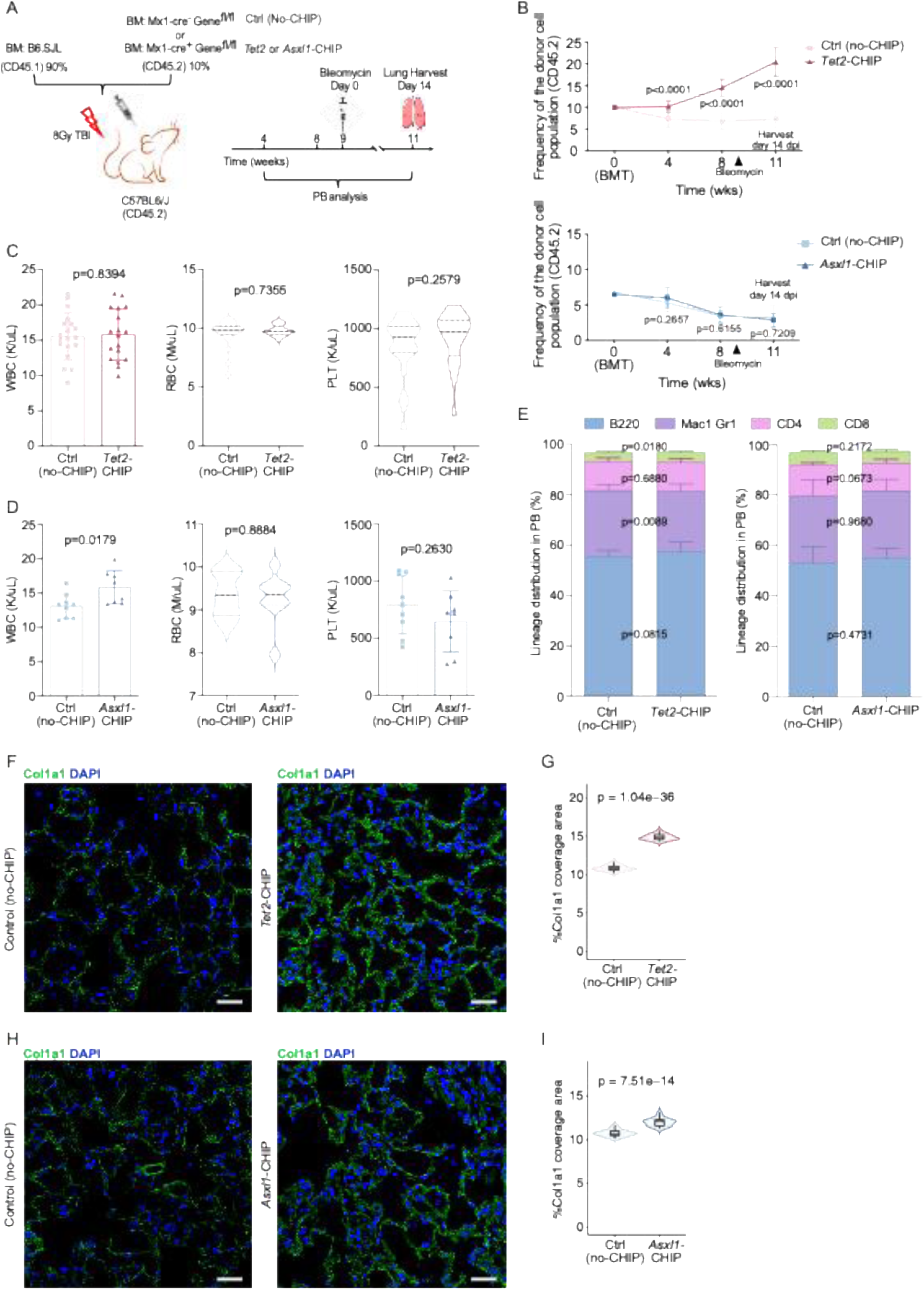
Hematopoietic characterization of the CHIP experimental models and lung collagen coverage area quantification post bleomycin administration. **a,** Schematic representation of the generation of the CHIP experimental models. **b,** Frequency of CD45.2^+^ cells in peripheral blood over time following bone marrow transplantation (BMT), demonstrating donor cell engraftment in the *Tet2*-CHIP and *Asxl1*-CHIP models. **c,d,** White blood cell (WBC), red blood cell (RBC) and platelet (PLT) count in peripheral blood at week 8 post–BM transplant in control versus *Tet2*- or *Asxl1*-CHIP mice. **e,** Peripheral blood (PB) lineage distribution at week 8 post–BM transplant in control versus *Tet2*- or *Asxl1*-CHIP mice. **f,h** Representative images of immunofluorescence (IF) staining for collagen type I alpha 1 (Col1a1) in the lungs of control and Tet2- or *Asxl1*-CHIP mice at 14 days post–bleomycin injury (14 d.p.i.). **g,i,** Quantification of Col1a1 coverage area in lung sections from control versus Tet2- or *Asxl1*-CHIPmice. Results represent independent experiments. n= 5-10 mice per genotype per experiment unless otherwise indicated. Data are presented as violin plots representing kernel density distributions, with the interquartile range indicated or as mean ± standard deviation (SD) for bar plots. Scale bars: 50μm.

**Extended data Fig. 4:**
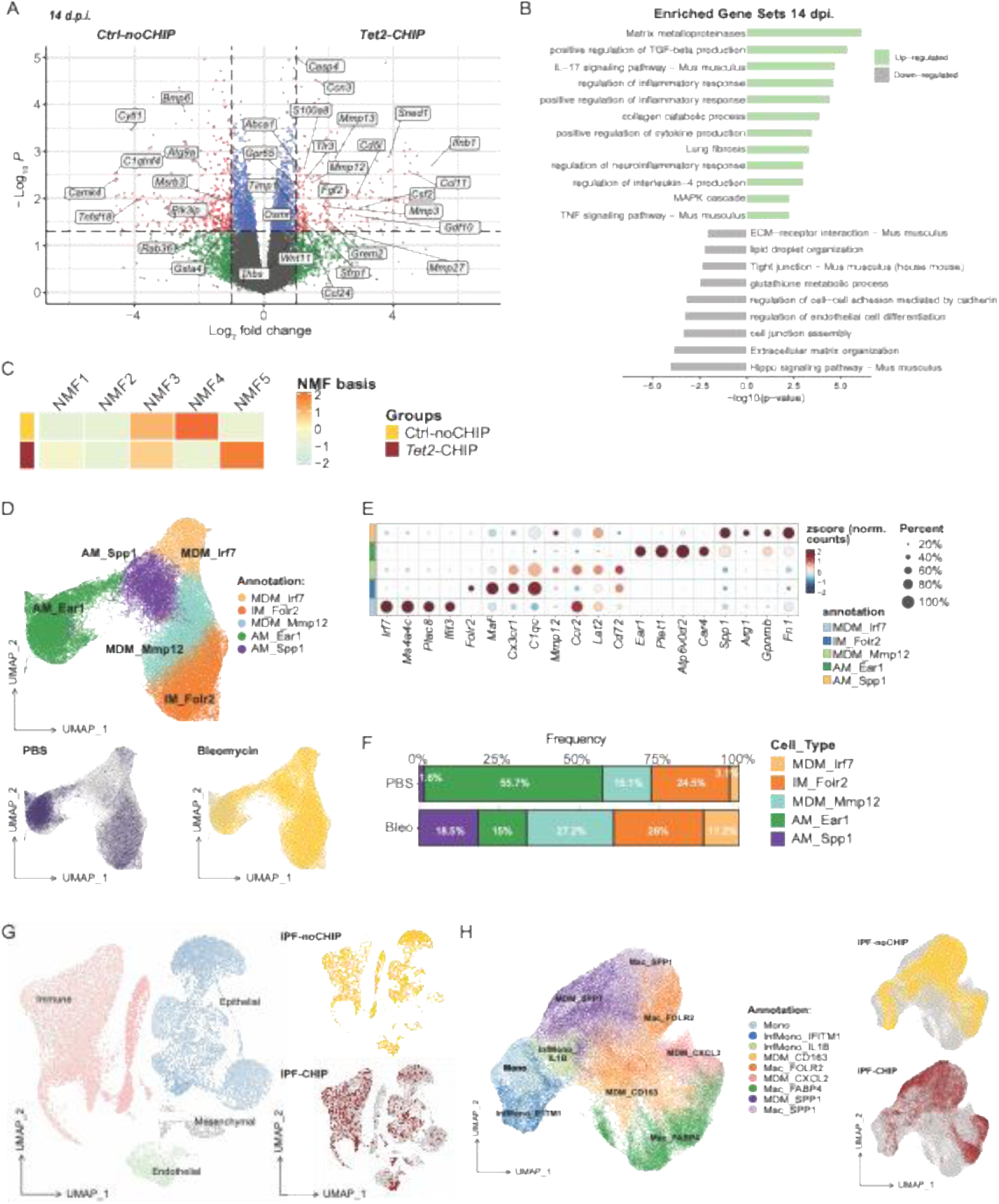
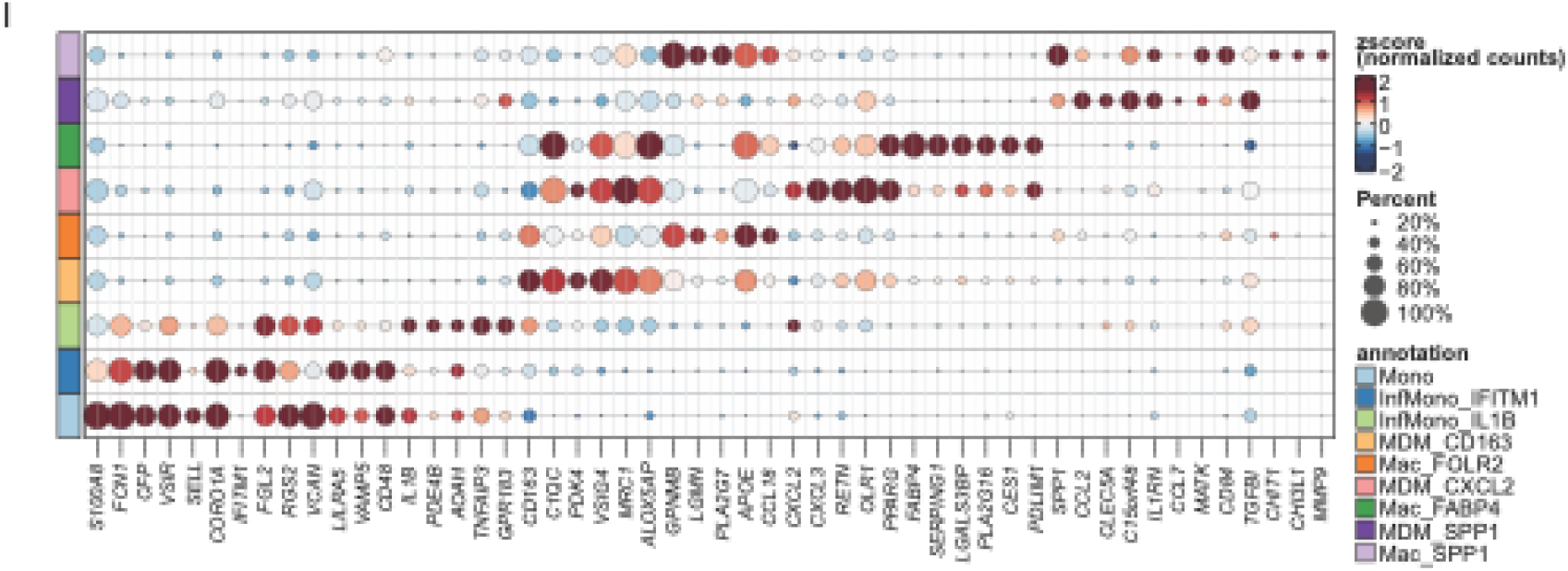
Transcriptional Signatures and Cellular Context of Macrophage Dysregulation in CHIP. **a,** Volcano plot showing DEGs between control-noCHIP and *Tet2-*CHIP mouse lung macrophages at 14 days post-injury (d.p.i.) from bulk RNA-seq. Genes with pvalue <0.05 and log2(fold change) > 1 are colored. **b,** Enriched gene sets identified via Over-Representation Analysis (ORA) analysis in *Tet2-*CHIP mouse lung macrophages at 14 d.p.i. **c,** Non-negative matrix factorization (NMF) of lung macrophage RNA-seq data at 14 d.p.i. showing two dominant metaprograms, NMF4 and NMF5, with preferential enrichment in control and *Tet2*-CHIP macrophages, respectively. **d,** UMAP plot showing macrophage clusters and treatment conditions in the public mouse scRNA-seq dataset (GSE280003) used for deconvolution analysis in Fig. 4d. **e,** Top marker genes of each cluster identified in panel **d** and used for the annotation. **f,** Quantification of cell type frequency for the clusters identified in panel **d** comparing untreated and bleomycin treated mice. **g,** UMAP plot showing cell lineage and CHIP status (IPF noCHIP or IPF-CHIP), identified by single-nucleotide variant (SNV) calling, in the human IPF scRNA-seq dataset (GSE227136) used for analysis in Fig. 4k-o. **h,** UMAP plot focusing on macrophage subclusters and their corresponding CHIP status. **i,** Top marker genes of each macrophage subcluster identified in panel h and used for the annotation.

**Extended data Fig. 5:**
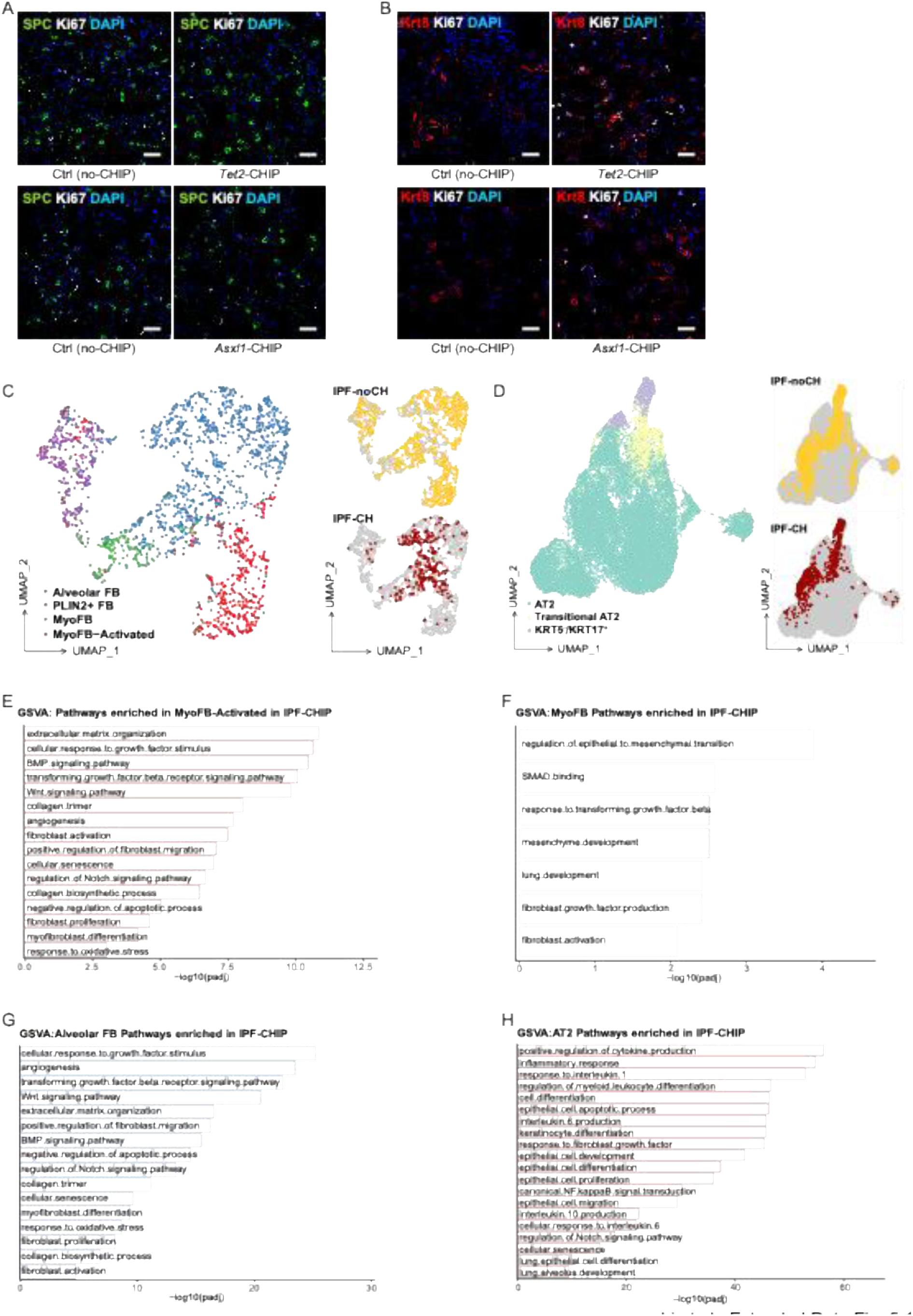
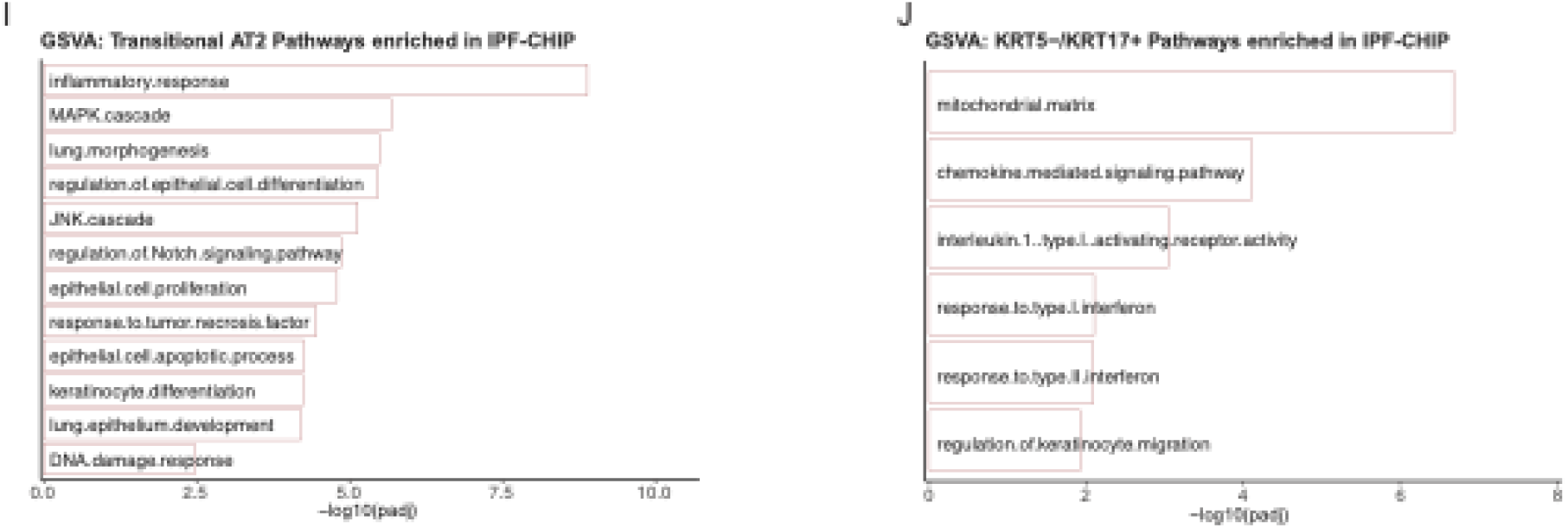
Transcriptional evaluation of mesenchymal and epithelial cell populations in IPF patients with and without predicted CHIP mutations. **a,b,** Representative IF images of proliferating Spc^+^ cells and Krt8^+^ cells assessed by Ki67 expression from Control-no-CHIP and *Asxl1-*CHIP. **c,d,** UMAP plot showing cell mesenchymal and epithelial lineage and CHIP status (IPF noCHIP or IPF-CHIP), identified by single-nucleotide variant (SNV) calling, in the human IPF scRNA-seq dataset (GSE227136) used for analysis in Fig. 5t**-u**. **e-g,** Enriched gene sets identified via Gene Set Variation Analysis (GSVA) analysis in activated myofibroblasts (**e**) myofibroblasts (**f**) and alveolar fibroblasts (**g**) of IPF patients with and without predicted CHIP mutations. **h-j,** Enriched gene sets identified via GSVA analysis in AT2 cells (**h**) Transitional AT2 cells (**i**) and KRT5-/Krt17+ aberrant epithelial cells (**j**) of IPF patients with and without predicted CHIP mutations. Scale bars: 50μm.

**Extended data Fig. 6:**
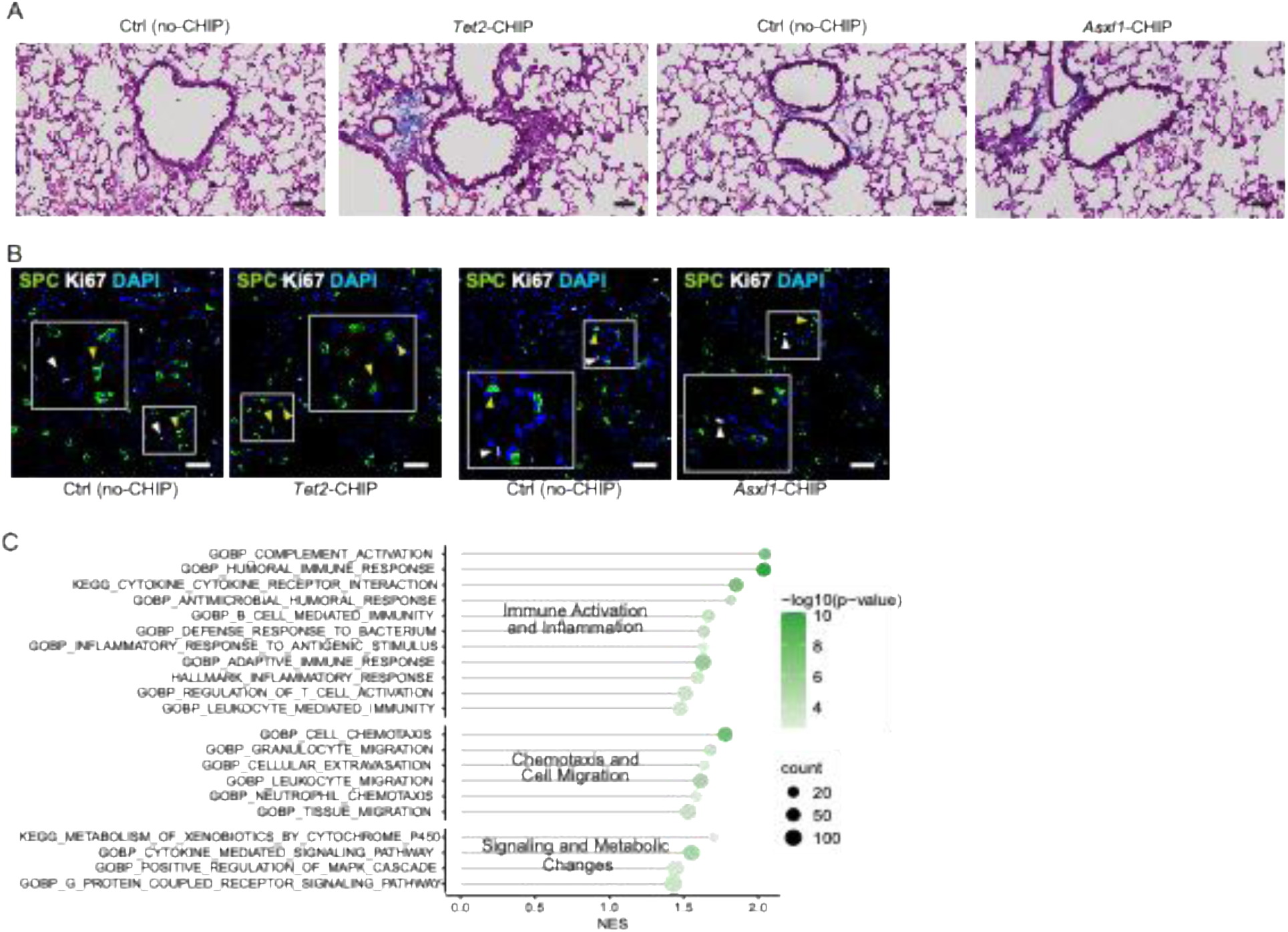
CHIP establishes a pre-injury parenchymal state marked by subtle structural and cellular changes. **a,** Representative lung histology images from Control-no-CHIP, *Tet2-* and *Asxl1-*CHIP stained with Masson’s Trichrome (collagen in blue). **b,** Representative IF images of Spc^+^ cell proliferation assayed by Ki67 co-localization in the lungs from Control-no-CHIP, *Tet2-* and *Asxl1-*CHIP in the absence of injury. **c.** Enriched gene sets identified via ORA analysis in *Tet2-*CHIP mouse lung macrophages in the absence of injury. Scale bars: 50μm.

